# Artificial Intelligence And First Principle Methods In Protein Redesign: A Marriage Of Convenience?

**DOI:** 10.1101/2025.05.12.653318

**Authors:** Damiano Cianferoni, David Vizarraga, Ana María Fernández-Escamilla, Ignacio Fita, Rahma Hamdani, Raul Reche, Javier Delgado, Luis Serrano

**Affiliations:** Centre for Genomic Regulation (CRG), The Barcelona Institute for Science and Technology, Dr. Aiguader 88, 08003 Barcelona, Spain; Universitat Pompeu Fabra (UPF), Barcelona, Spain; ICREA, Pg. Lluis Companys 23, 08010 Barcelona, Spain; Instituto de Investigación, Desarrollo e Innovación en Biotecnología Sanitaria de Elche (IDiBE), Universitas Miguel Hernández, 03202 Elche, Spain; Universitat de Barcelona (UB), Barcelona, Spain; Instituto de Biología Molecular de Barcelona (IBMB-CSIC), Parc Científic de Barcelona, Barcelona, Spain

**Keywords:** protein design, artificial intelligence, force field, crystallographic structure

## Abstract

Since AlphaFold2’s rise, many deep learning methods for protein design have emerged. Here, we validate widely used and recognized tools, compare them with first-principle methods, and explore their combinations, focusing on their effectiveness in protein redesign and potential for therapeutic repurposing. We address two challenges: evaluating tools and combinations ability to detect the effects of multiple concurrent mutations in protein variants, and leveraging large-scale datasets to compare modeling-free methods, namely force fields, which handle point mutations well with limited backbone rearrangement, and inverse folding tools, which excel at native sequence recovery but may struggle with non-natural proteins. Debuting TriCombine, a tool that identifies residue triangles in input structures, matches them to a structural database, and scores mutants based on substitution frequencies, we shortlisted candidates, modeled them with FoldX, and generated 16 SH3 mutants carrying up to 9 concurrent substitutions. The dataset was expanded to include 36 mutants and 11 crystal structures (7 newly solved), along with a parallel set of multiple non-concurrent mutants from three additional proteins. For broader validation, we analyzed 160,000 four-site GB1 mutants and 163,555 (single and double) variants across 179 natural and de novo domains. We show that combining AI-based modeling tools with force field scoring functions yields the most reliable results. Inverse folding tools perform very well but lose accuracy on less-represented proteins. First-principle force fields like FoldX remain the most accurate for point mutations. All methods perform worse when applied to unsolved de novo models, underscoring the need for hybrid strategies in robust protein design.

## INTRODUCTION

Until recently, protein design was typically based on the use of energy functions derived from first principles (Rosetta^1^, FoldX^2,3^, Gromacs^4^, I-TASSER^4,5^), as well as side chain and backbone exploration strategies. Most challenges were related to the exponential size increase of sequence space, the consequential combinatorial problem and with trapping into local minima while assigning rotamers sequentially^6,1^.

Recent advancements in machine learning^7,8,9^ have fueled the potential of de novo design, which has been demonstrated to be highly effective in generating novel molecules with desired functions^10,11^. Deep-learning based tools have been engineered to approach the problem of finding a sequence for a backbone structure (*ProteinMPNN*^12^, *Esm_inverse*^13^) by learning the protein language, offering an alternative route to the explicit extraction of evolutionary information from multiple sequence alignments^14^. Other approaches making use of diffusion models can generate accurate backbones very similar to experimental structure^15,16^. Despite the promising novelty and impressive outcomes, the use of de novo solutions^17^ in the protein therapeutics market remains scarce. This market predominantly favors established molecule variants, exemplified by antibodies, which are gradually being integrated alongside FDA and EMA^61^.

Although deep learning tools for predicting protein variants show strong capabilities, they are often validated on large mutant datasets to assess their predictive power. A key issue is that these datasets often report fitness metrics influenced by many factors beyond protein stability that are hard to disentangle. Additionally, they typically feature only one or two mutations per domain^18^. These tools are often evaluated by how well they predict existing protein variants, rather than by identifying the best de novo designs with minimal attempts. While validation on large datasets of single and double mutants provides useful insights into how broadly a model can generalize across sequences with very limited Hamming distance (few mutations), it does not fully capture the challenges of multi-site redesign. In therapeutic design scenarios, it is not enough for a model to capture general trends. Instead, the model must reliably predict the effects of variants with multiple mutations, accurately model the resulting structural changes, and be evaluated using meaningful functional and biochemical stability metrics.

Our study focuses on structural design in the context of protein redesign involving multiple mutations affecting interacting residues. While the term “de novo” design is often overused in the field, we believe it is inappropriate when the protein’s sequence or structure remains related to a natural counterpart. In such cases, it is more accurate to refer to these proteins as redesigned, especially when they retain a recognizable fold. Our goal is not to provide a comprehensive comparison of all available methods; instead, we focus on those that are widely validated, commonly used by the community, and capable of handling multiple mutations. To achieve this, we selected two distinct scenarios. In the first, we assess the efficacy of various methods in designing many-mutation variants by using biochemical stabilities of mutants and wild types. We included redesigned variants of the entire hydrophobic core of an SH3 domain, as well as three sets of mutants from the literature, featuring multiple non-interfering mutations in larger proteins. In the second scenario, we evaluate the performance of inverse folding tools in predicting stable variants between extensive mutation datasets involving few residue positions: one case involves mapping the full mutation landscape for four residues^19^, while the other involves single and double mutations across numerous natural and artificial domains^20^.

For the first challenge we have assembled a dataset of 36 multiple mutants (ranging from 3 to 9 residues) of the spectrin SH3 domain’s hydrophobic core (9 residues) with precise chemical denaturation stability measurements^21^. Sixteen mutants were generated using our new and simple approach: *TriCombine.* This approach matches, within a user-defined tolerance, triangles of residues from an input backbone to a structural database of triangles (TriXDB) and scores protein variants based on substitution frequency (see Methods and Supplementary Figure 1). Designed to work with *FoldX*, *TriCombine* is part of the ModelX toolsuite^32,33,34^. It streamlines the sequence search for a backbone to identify variants that preserve the wild type structure while introducing small backbone conformational diversity. This approach was developed to address the above-mentioned limitations of traditional rotamer-based protein design by simplifying sequence search on a given protein backbone. Furthermore, we determined the structure of seven of the designed SH3 multiple mutants. We used the stability measurements and crystal structures (the seven determined here plus four previously published ones^21,22^ to compare the most relevant protein design tools and determine the most accurate methods for approximating both structure and stability. Tested tools and combinations^23^ fell into three categories: structure prediction from protein sequence (*EsmFold*^14^, *AlphaFold2*^24^, *RoseTTAFold*^25^); sequence prediction from a backbone of input (*Esm_inverse*^26^, *proteinMPNN*^27^, *TriCombine*) and force fields based (*Rosetta*^27^, *FoldX*^2^). We further supplemented the analysis with a dataset of three groups of mutants counting 15 variants from literature derived from human acetylcholinesterase (hAChE)^28^, bacterial phosphotriesterase (PTE)^28^, and fibronectin type III domain of the protein tenascin (TNfn3) variants^29^.

For the second challenge, we investigated the fitness landscape of mutants involving four interacting residues within the Protein G domain B1 (GB1), two of which interact with the IgG partner^19^. In addition, we curated a dataset of 163’555 free energy variations for single and double mutants divided in two subsets of 155 artificial domains and 58 natural ones^20^. We compared the fitness experimental data with sequence predictions derived from inverse folding tools (*Esm_inverse*^26^, *ProteinMPNN*^12^), fixed backbone forcefield tools (*FoldX*^30^ and *Rosetta* ^1^) and our new tool *TriCombine* to test its performances as a scorer beyond its original function to generate mutants.

## RESULTS

### New Independently Designed SH3 Poly-Mutant Variants Using the TriCombine Algorithm

In our initial challenge, we chose a protein design task that seemed relatively straightforward: the redesign of the hydrophobic core of a small SH3 spectrin domain, which consists of 62 residues. This core includes nine residues with a solvent accessibility of less than 10%^31^. These residues are Val 9, Ala 11, Val 23, Met 25, Leu 31, Val 44, Val 46, Val 53 and Val 58 (PDB ID: 1shg). We also used an expanded and strained spectrin SH3 hydrophobic core variant (PDB ID: 1e6g)^21^ to have a different design starting point. In this structure the core residues are Val 9, Val 11, Leu 23, Ile 25, Leu 31, Val 44, Val 46, Ile 53 and Leu 58 (Fig. 1A).

**Fig. 1.**
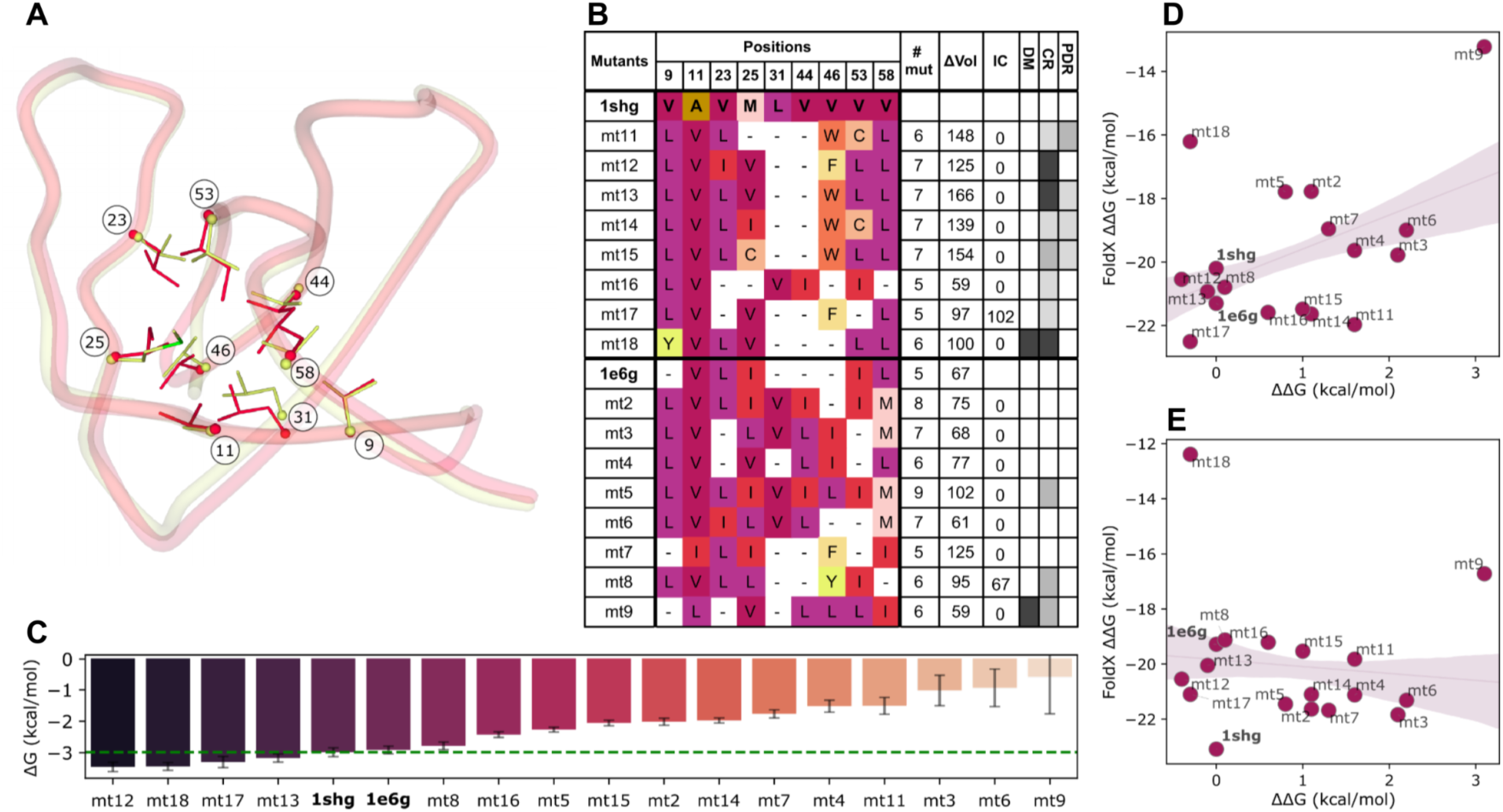
A) In red, tube visualization of 1e6g backbone showing the 9 residues of the hydrophobic core as sticks. In yellow, tube backbone of 1shg showing hydrophobic core residues as sticks. B) Table shows the 16 candidates core mutations, compared with the two starting structures sequences. A “-” symbol indicates the same amino acid as in the WT structure (1shg). Table columns from left to right: name of the mutant, identities of the mutated residues at a certain position, number of mutations for the mutant (# mut), predicted variation in Å^3^ for the hydrophobic core (ΔVol), total cavity volume generated in Å^3^ (IC), whether the mutant has total energy sensibly worse than its reference structure (DM), presence of residues with Van der Waals clashes (CR), residues with polar atoms desolvated not making a H-bond (PDR). Light gray indicates at least one core residue with a ΔΔG with respect to its reference structure between 0.5 and 0.8 kcal/mol. Dark gray shows more than one residue between 0.5 and 0.8 kcal/mol or at least one higher than 0.8 kcal/mol. Black boxes indicate more than one core residue above 0.8 kcal/mol. C) Experimental ΔG of folding for the 16 mutants, 1shg and 1e6g variants. A green dotted line marks WT stability. D) Comparison of experimental ΔΔG with FoldX energy predictions for mutants modeled over 1shg template previously mutated to Alanine in the hydrophobic core positions. E) Comparison of experimental ΔΔG values with the FoldX energy predictions for mutants modeled over the 1e6g template previously mutated to Alanine in core positions.

Starting from these two models, we redesigned 16 variants using *TriCombine*, eight for each model. *TriCombine’*s approach follows the ModelX philosophy, formerly shown in DNA^32^, RNA^33^ and protein-protein docking^34^ that assumes that the landscape of interacting residues captured in high resolution crystals from the Protein Data Bank^35^ (PDB) is huge but limited. Based on these considerations, we developed a novel ModelX database (TriXDB) containing all possible combinations of three residues that contact each other in naturally observed conformations (see Methods and Supplementary Fig. 1). These three residue groups enable the simultaneous placement of multiple side chains that are compatible with the input backbone (*TriCombine*).

The SH3 variants were generated by mutating to Alanine the nine core residues before running *TriCombine*, eliminating in this way any bias imposed by rotamer placement constraints due to the WT side chains. As we redesigned the hydrophobic core, we excluded charged residues (Asp, Glu, Lys, Arg), large polar residues (Asn, Gln), and Glycine from the final ranking process, so that no combinations with such amino acids were scored (see Methods). We included Histidine, as it can form favorable stacking interactions with aromatic residues, and we retrieved only TriXs that had at least two internal side chain contacts. To account for potential small backbone shifts upon redesign, we allowed for 0.3Å tolerance (‘*dubiety’* parameter) in superimposition between TriX’s Cα and Cβ atoms and the anchoring residues in the SH3 structure (as shown in Supplementary Fig. 1B and explained in Methods). Finally, TriXs from PDB structures with >30% sequence similarity to the SH3 domain were excluded, to demonstrate the method’s independence from prior SH3 core information. Scores for the top 500,000 combinations of both design batches (1shg, 1e6g) are stored separately in two relative output files. The top 10,000 ranking sequences for each chosen SH3 starting structure (1shg and 1e6g) were modeled over their corresponding input PDB using *FoldX* BuildModel, also allowing the replacement of a maximum of 2 clashing residue substitutions by utilizing the *FoldX* PSSM command to explore point mutations stabilizing the models (see Methods).

The sixteen candidates were chosen for experimental validation based on the following criteria: i) Candidates free of structural issues within the backbone of input (mt2, mt4, mt6, mt7, mt16, mt17). ii) Candidates potentially affected by minor structural issues (mt5, mt11, mt12, mt13, mt14, mt15). iii) Candidates with strong structural issues (mt9, mt18). We considered structural issues: Van der Waalś clashes, cavities, hydrophobic desolvation and unsatisfied polar groups (see Fig. 1B, Methods and Supplementary Table 1). All selected sequences should have at least 5 out of the 9 core residues mutated.

Another filter for candidate selection was put in place to ensure the suggested combinations of amino acid were not previously observed in nature: a sequence search within the SH3 domain family was performed and each of the 16 mutants were aligned with the “full family” alignment for SH3_1 Pfam^36^ family (ID:PF00018). Our analysis revealed that none of the core residue combinations generated in our study matched those previously reported. Out of the 16 identified variants, those with sequences most similar to any of the 92,969 known ones shared 7 out of 9 amino acids (mt7 and mt8), while the one least similar shared only 5 out of the total 9 amino acids (Supplementary Note 2 and Supplementary Table 2).

We expressed, purified and determined the folding stability for the 16 mutants, the WT (1shg) and the expanded variant (1e6g). Five candidates resulted equally or more stable than the WT (mt8, mt12, mt13, mt17, mt18), six were less stable but folded (mt2, mt5, mt7, mt14, mt15, mt16), two were partially unfolded (mt4, mt11) and three of them mainly unfolded (mt3, mt6, mt9) (Fig. 1C and Supplementary Fig. 2).

Most mutants designed over 1shg structure were found folded (mt12, mt13, mt14, mt15, mt16, mt17, mt18), while those based on the swollen and strained 1e6g structure were generally less stable (mt2, mt5, mt7), partially unfolded (mt4) or unfolded (mt3, mt6, mt9). Surprisingly mt8, which was designed over 1e6g having a cavity, was found to be significantly more stable than expected, indicating that the structure transitioned from a strained state to a stable one as the cavity compacted reducing the final volume (Figure 1B; see below in structural prediction).

We then modeled all designs with *FoldX* using both 1shg (Fig. 1D) and 1e6g (Fig. 1E) structures and compared the predicted stability with experimental values. This analysis revealed that we got reasonable predictions when using the WT structure as template, but not when using the core-expanded 1e6g structure. This limitation also explains why mt8, designed with a cavity on 1e6g, turned out to be stable. As shown in a recent *FoldX* update^37^, the tool is not well suited to model mutations in strained proteins, such as those engineered by introducing large residues into the core, as in the case of 1e6g (Figure 1B; see ΔVol relative to the wild type 1shg), where subsequent mutations to smaller residues relieve the strain.

### Comparing Predictions with Biochemically Measured Stabilities for Extensively Redesigned Variants

We expanded our SH3 mutant dataset by adding stability measurements from 20 SH3 hydrophobic core variants previously described by Ventura et al. ^21^ bringing the total to 36 experimental ΔG measures. We then conducted a correlation analysis between experimental stabilities versus the predicted scores generated by the following tools and their combinations: *TriCombine*, *FoldX*, *Rosetta*, *EsmFold*, *RoseTTAFold*, *Esm_inverse*, *ProteinMPNN* and *AlphaFold2* (see Methods for details). In total we had 20 metrics that can be divided in four categories: I) scoring functions based on fixed input backbone (*TriCombine*, *Esm_inverse*, *proteinMPNN*, *FoldX*, *Rosetta*) using the wild type (WT) SH3 as a reference; II) AI tools that generate and assign a reliability score to a structural model starting from a protein sequence (*EsmFold*, *RoseTTAFold*, *AlphaFold2*); III) Scores obtained from the previous AI generated models evaluated with *FoldX* (*EsmFold_F*, *RoseTTAFold_F*, *AlphaFold2_F*) and *Rosetta* force fields (*EsmFold_R*, *RoseTTAFold_R*, *AlphaFold2_R*); IV) Scores from models initially generated by the tools in group II, then subjected to relaxation (see Methods) and finally scored using either *FoldX* (*EsmFold_r_F, RoseTTAFold_r_F, AlphaFold2_r_F*) or *Rosetta* (*EsmFold_r_R, RoseTTAFold_r_R, AlphaFold2_r_R*), respectively. For all the metrics and all datasets we compared variants predictions with the one produced for the wild type variant.

We measured the number of true positive predictions (TP), representing variants predicted with a superior score compared to the one predicted for the WT and confirmed experimentally to be more stable. As well as we measured the true negatives (TN), false positives (FP), and false negatives (FN) in this respect. With that, we assessed methods and their combinations using balanced accuracy^38^, aiming to rank them based on their ability to suggest variants superior to the wild type. We also determine the correlation between the experimental ΔΔG values and the predicted scores of the different methods.

In Figure 2A, for the SH3 dataset, we see *EsmFold_F* and *AlphaFold_R*, two modeling methods coupled with *FoldX* or *Rosetta* force field functions to be the ones with best accuracy. As expected, fixed-backbone, force-field-based methods (*FoldX* and *Rosetta*) show limitations with SH3 case accuracies, since the selected variants were intentionally designed with defects that would induce backbone rearrangements (Figure 1B). *Esm_inverse* and *ProteinMPNN* also struggled to identify mutants surpassing the WT. However, as depicted in Supplementary Fig. 5, this is due to the overestimation of the WT that is our unique reference in a redesign experiment. *EsmFold* showed the strongest Pearson correlation with experimental data (Fig. 2A and Supplementary Fig. 5). It also demonstrated a high proportion of TN predictions, albeit with limited TPs. Additionally, it performed well with the *FoldX* force field (*EsmFold_F*) but not so well with *Rosetta’s* (*EsmFold_R*). *RoseTTAFold* works well when coupled with *Rosetta’s* force field (*RoseTTAFold_R*).

**Fig. 2.**
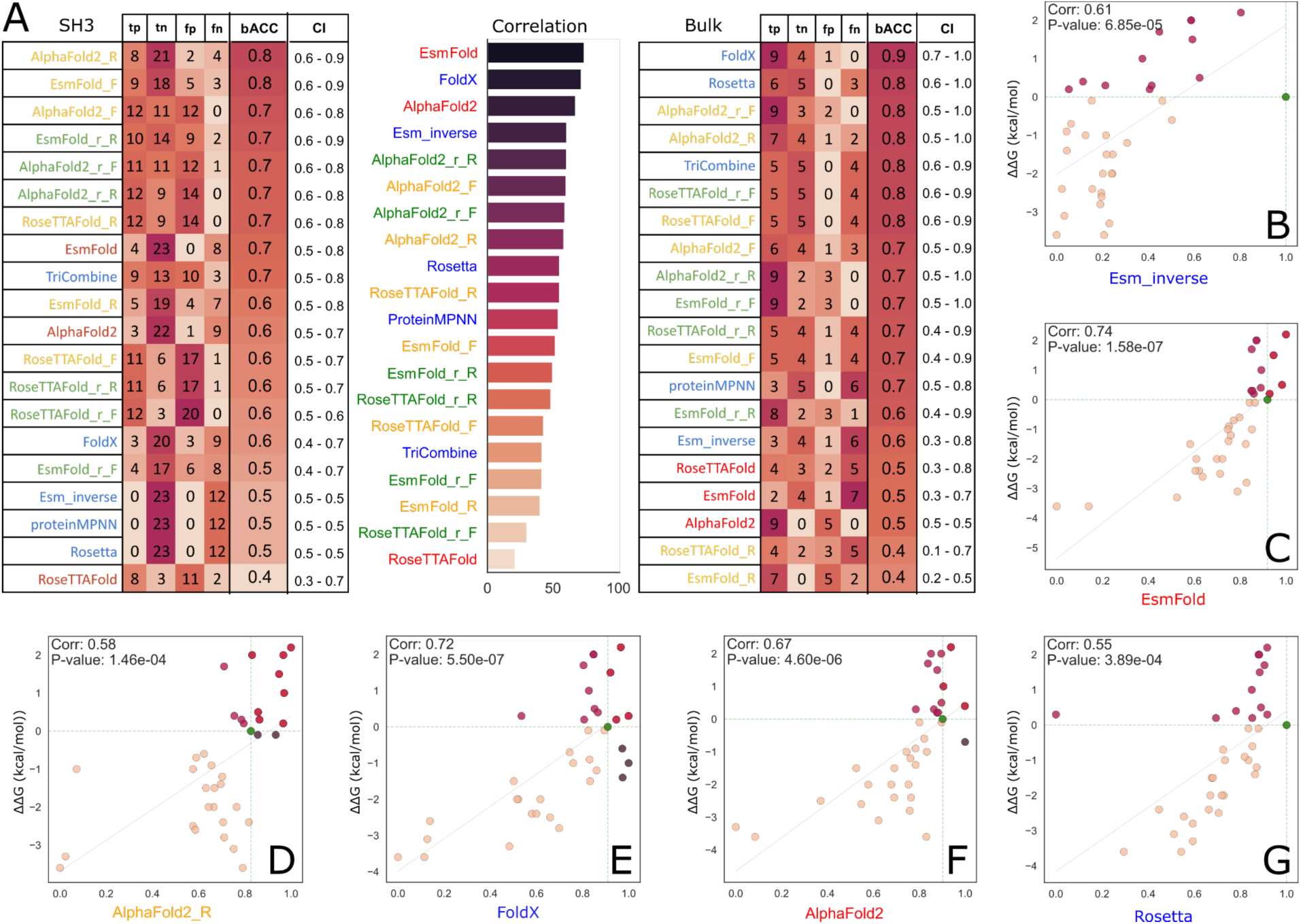
A) Table showing true positive (tp), true negative (tn), false positive (fp) false negative (fn) and balanced accuracy (bACC) with confidence intervals (CI) of foldability prediction for all tested approaches over the 36 stability data points for SH3 variants (left). In the middle, a histogram compares the correlations of predictions with experimental ΔΔGs on a percentage scale. On the right the same comparison shown on the left but for the literature batch. B-G) scatterplot of selected scorers showing experimental ΔΔGs versus predicted Δ-scores. Dotted green line shows WT values on the two axes, bottom left point shows variants correctly predicted as less stable than the WT, top right are correctly predicted as more stable. Bottom right shows overpredicted variants and top left underpredicted ones. Finally in green the WT variant point.

We expanded our analysis by incorporating a “bulk” dataset, composed of three additional groups of mutants from the literature (Figure 2A). The proteins involved are: human acetylcholinesterase (hAChE)^28^, a bacterial phosphotriesterase (PTE) variants^28^, and fibronectin type III domain of the protein tenascin (TNfn3)^29^. These datasets include a substantial number of mutations (17, 30, 42, 51 and 67 for hAChE; 3, 9, 19 and 28 for PTE; 14, 7, 22, 9, 19 and 12 for TNfn3). While the mutations are numerous, they are mostly independent of each other. Because these published mutants were mainly selected for increased stability, the dataset is relatively unbalanced, with fewer destabilized variants in the analysis. Stability was measured through melting or inactivation temperatures, meaning that ΔG cannot be inferred in a way that allows direct comparison across different proteins, so we relied only on balanced accuracy to rank the methods in this dataset.

Here, we found that force-field-based predictors *FoldX* and *Rosetta* perform best, most likely because the mutations affect residues not interacting with each other, they are most likely stabilizing, and therefore they will not induce conformational changes. *AlphaFold2*, as in the previous multi-mutant analysis, exhibits strong predictive capability when combined with *FoldX* or *Rosetta*. *TriCombine* maintains consistently average performances, showing relative independence from mutation type, probably thanks to its ability to envision minor backbone adjustments that allowed uncertainties in the superimposition of the selected TriXs.

### Sub-Amstrong structural prediction of SH3 variants

We then attempted the crystallization of ten redesigned SH3 variants (mt4, mt5, mt6, mt8, mt12, mt13, mt15, mt16, mt17, and mt18), which included those with the most unexpected stabilities with respect to *FoldX* predictions and successfully obtained structures for seven of them: mt8, mt12, mt13, mt15, mt16, mt17, and mt18. In general, the resolution of the structures was excellent (the worst was less than 1.4Å), the observed conformational changes were local and small (Supplementary Fig. 3A), as the maximum measured Cα Root Mean Square Deviation (RMSD) between the mutants and their respective starting backbone structure (1shg and 1e6g) was 1.0Å after structural alignment (Supplementary Fig. 3B).

Among the crystallized mutants, mt16 was designed to be mostly free of structural issues and showed the lowest Cα Root Mean Square Deviation (RMSD) with respect to 1shg. The same was found also for mt12, having two residues with moderate Van der Waals clashes, as well as for mt17 (Supplementary Fig. 3B,C). Mutant mt18, designed with significant van der Waals clashes, was more stable than the WT and had larger overall backbone deviations (Supplementary Fig. 3B). Mutant mt8, designed on 1e6g and expected to have a large cavity, packed it back and approximated its backbone to that of the reference WT structure (1shg) thus supporting the idea that the reference structure 1e6g was strained. All mutants having an aromatic residue at position 46 (mt8 a Tyr, mt12 and m17 a Phe, mt13 and mt15 a Trp) were predicted to desolvate a CO group from residue 24 backbone to different extents. In all cases we saw a displacement of the main chain around residue 46, allowing the CO group to have access to the solvent (Supplementary Fig. 3E). We discovered two alternative conformations of Trp 46 for crystals of mt13 and mt15. In mt13, the Trp 46 conformations consist of flipped poses of the amino acid side chain, which overlap at the Cα and Cβ positions. In mt15, the conformational change is more evident and involves the flipping of the introduced Cys 25. In one case, Cys 25 frees up the core, creating a cavity for interaction with the Trp that packs against the core. In the other case, Cys 25 fills the core space, and Trp 46 flips towards the outside (Supplementary Fig. 4)

Then we assessed which modeling tool produced structures that most closely matched the 7 SH3 crystals determined here (mt12, mt13, mt15, mt16, mt17, mt18, and mt8), three more multiple mutants previously determined (1e6g, 1e6h, 1h8k) and the WT (1shg). We evaluated the following methods capable of generating 3D models from scratch using only the sequence: *EsmFold*, *AlphaFold2*, *RoseTTAFold*. We utilized models generated by repairing the previous ones with *FoldX* RepairPDB (“*EsmFold_r_F*”, “*AlphaFold2_r_F*”, and “*RoseTTAFold_r_F*”), and with Rosetta’s relax routine (“*EsmFold_r_R*”, “*AlphaFold2_r_R*”, “*RoseTTAFold_r_R*”). Finally, we included fixed-backbone side-chain design methods by *FoldX* and *Rosetta*.

To ensure a reliable structural comparison among models, we standardized all structures by including only residues from Lys 6 to Asp 57. We removed water molecules and added hydrogen atoms. Subsequently, all models were structurally aligned using Mustang^39^ against the relative crystallographic structures. Global Distance Test (GDT) scores were then measured using all atoms and thresholds of 0.5Å and 1Å (Fig. 3A and Supplementary Table 3). Mean values were calculated for each metric from the 11 measurements of the 11 mutant structures. We also analyzed single side chain atomic RMSD relative to crystal structures, considering both alpha carbon and all side chain atoms (Supplementary Table 4). In one analysis, we averaged the RMSD per mutant for each method (Fig. 3B and Supplementary Fig. 6A), while Supplementary Figure 6B shows the complementary analysis where we averaged across methods to highlight the performance per residue per mutant.

**Fig. 3.**
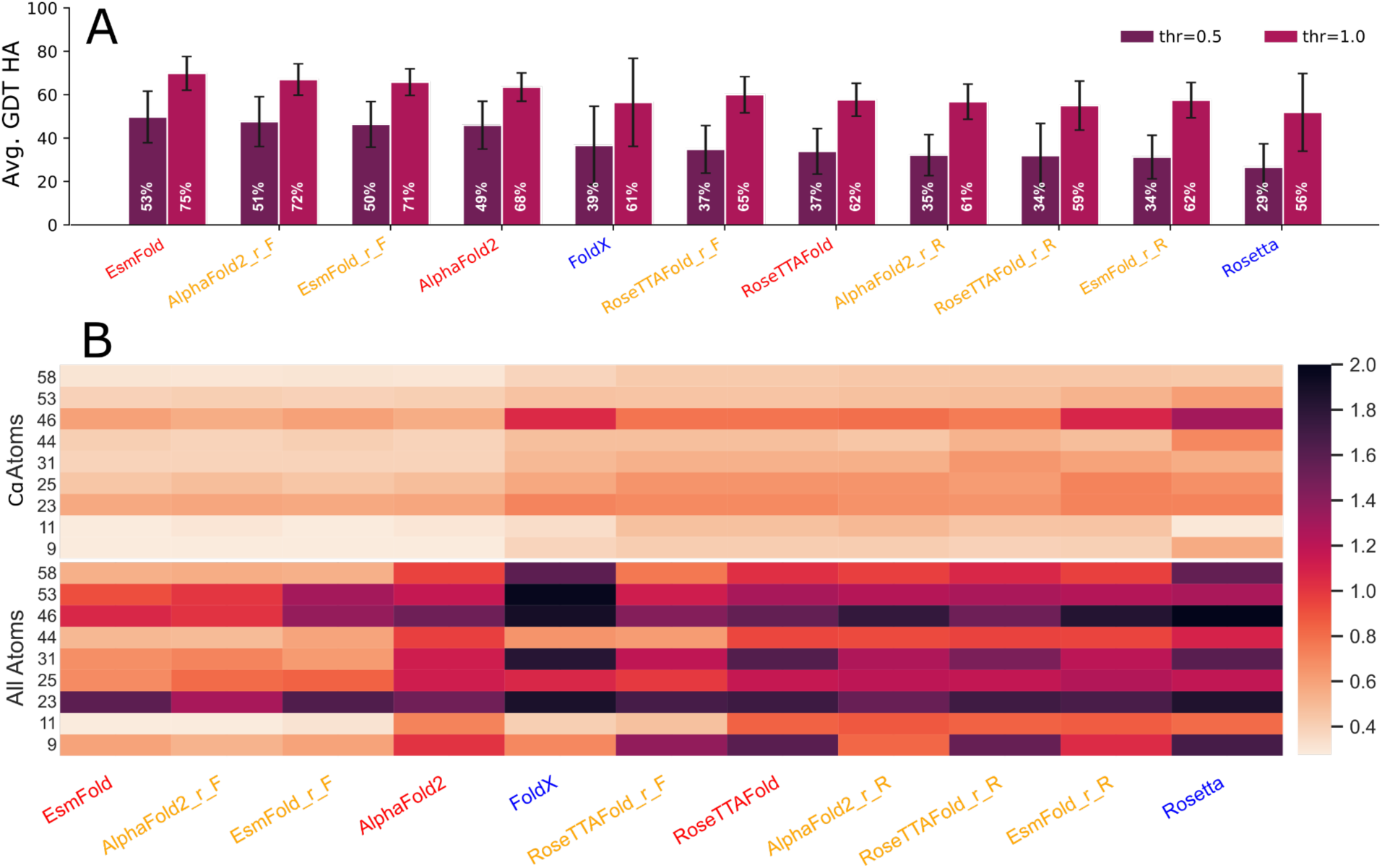
A) Barplot illustrating average GDT scores across 11 crystallographic structures of SH3 domain mutants. Darker bars show the percentage of atoms predicted within 0.5 Å from expected positions; Clearer bars: within 1Å. Tool names are color-coded: red for those modeling starting from sequence, orange for the same ones but later relaxed with FoldX (F) or Rosetta (R), blue for fixed backbone ones using 1shg as template. B) Heatmaps showing hydrophobic core side chain RMSD for each modelling method averaged over 11 crystallized variants. Above the average Cα RMSD, below the whole-atom one. Values were capped to an upper limit of 2Å. The complete RMSD analysis is available at Supplementary Fig. 6 and Supplementary Table 4.

Without strong significance, the analysis ranked *EsmFold* with the highest accuracy, followed by *AlphaFold2* models repaired with the *FoldX* force field (*AlphaFold_r_F*), a trend also observed for *RoseTTAFold_r_F*. In contrast, *<u>RoseTTAFold</u>* and methods involving Rosetta backbone relaxation performed worse than directly modeling side chains on the wild type backbone with *FoldX* (Fig. 3A). *FoldX* was particularly effective at placing rotamers where the backbone was accurately modeled, as seen at residues 9, 11, and 44 (Fig. 3B). *EsmFold* generally predicted both Cα positions and side chains well; however, when the Cα atom was slightly misplaced, *FoldX* repair often worsened the side chain (residues 46, 53, 25; Fig. 3B), suggesting that *EsmFold* could still infer the correct rotamer orientation despite the displaced Cα, while *FoldX* optimized the rotamer for an incorrect backbone. In contrast, *AlphaFold2* predicted Cα positions as accurately as *EsmFold* but produced less accurate rotamers, its side chains often improved after refinement with *FoldX* (Fig. 3B). These results, consistent with our balanced accuracy analysis, highlight the benefit of using modeling tools for structure generation and directly applying force fields for evaluation, skipping further structural refinement. In the case of *EsmFold*, the best predictions are achieved by using the original, unrefined models while still relying on force fields for scoring. Thus, refinement is unnecessary unless detailed inspection of the three-dimensional structure is required.

### Prediction of Large Scale Few-Mutation Stable Variants Using Modeling-Free tools

To further validate our observations, we showed the comparison in performance between two classes of predictors all working with the same input backbone structure. On one hand we tested the two most relevant inverse folding algorithms, *ProteinMPNN*^12^ and *Esm_inverse*^26^, that showed approximately 50% of sequence recovery on native protein backbones. On the flip side, two established force field-based methods (*FoldX* and *Rosetta*) using the same backbone of input, while we have also integrated our novel tool, *TriCombine*.

To test the different methods, we selected three different datasets. The first one comprises nearly 160,000 fitness points derived from non-consecutive mutated residues in contact between each other (Val 39, Asp 40, Gly 41 and Val 54) of a Protein G domain B1 (GB1) as shown in Figure 4A. Each variant’s fitness value was determined by measuring the ratio of counts for the mutant after and before mRNA affinity selection (binding IgG-Fc partner) normalized against the WT variant^19^. To generate the predictions we used a crystallographic model of the C2 fragment of streptococcal protein G in complex with the Fc domain of human IgG (PDB id: 1fcc) differing on one residue position (Ile 7) from the one used to generate the dataset^19^ (Leu 7). Leu 7 is peripheral residue, not part of the mutation sites that was changed to leucine with *FoldX* in 1fcc, showing a ΔΔG below the significance (0.8 kcal/mol) of 0.4 kcal/mol, confirming its neutral impact and the use of 1fcc structure for our purposes. The second and third datasets were derived from the Mega-scale experimental analysis of protein folding stability in biology and design^20^ that comprises almost one Million cDNA display proteolysis stability points that we filtered to nearly 170’000 high quality measurements (Figure 4B) (see Supp. Material for details). The second dataset contains point and double mutations on natural proteins for which a high-resolution crystallographic structure is available. The third one contains mutations on artificial proteins, for which no structural data were available. Double mutants, in both cases, were restricted to those involving residues in contact with each other.

**Fig. 4.**
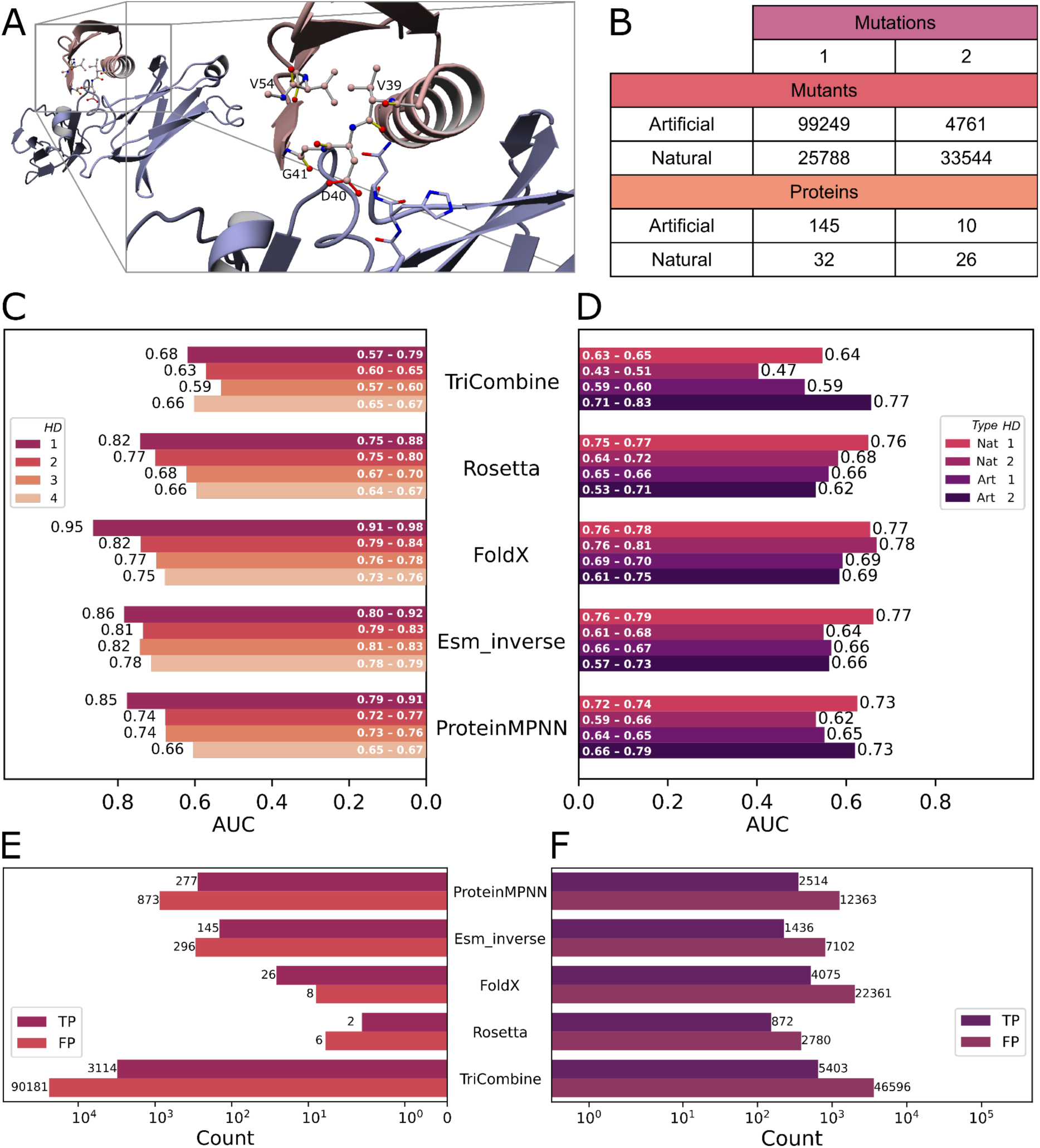
A) Wild type GB1 structure (pink) bound to IgG ligand (purple), with the mutated residue highlighted. The right panel shows a close-up of the four residues and two interacting ligand residues (Asn and His, purple). B) Diversity of mutants from the filtered Mega-scale dataset. C) Histogram comparing AUC performance of five fixed-backbone predictors for 1-4 Hamming Distance mutants in the GB1 dataset. D) Histogram comparing AUC performance for 1-2 Hamming Distance mutants from Natural and Artificial proteins in the filtered Mega-scale dataset. E) True Positive (TP) and False Positive (FP) counts for the five predictors, relative to experimental ΔΔG > 0 against wild type in the GB1 dataset. F) TP and FP count for the five predictors, comparing experimental ΔΔG > 0 against wild type in the filtered Mega-scale dataset.

*Esm_inverse* was launched employing a specific routine to extract sequence scores, similarly for *ProteinMPNN*. *FoldX* buildmodel was used to mutate the four residues for each combination and measure the correspondent stability ΔΔG of the complex. The *Rosetta* FastDesign routine was employed with the fix backbone option to obtain scores for the variants, additionally, *TriCombine* scores were included in the analysis (see Methods for details). We then analyzed the results to determine which tool identified the most true positive variants. Similarly to the SH3 case, true positives were defined as those with equal or better experimental fitness compared to the WT. We stratified the results by the number of mutations (ranging from 1 to 4) and assessed overall performance by generating ROC curves. For each group, we calculated the Area Under the ROC Curve (AUROC) and estimated 95% confidence intervals using bootstrap resampling, for each protein, we set the threshold for calling positives and negatives to the wild type’s values.

For the GB1 dataset we found that for single point mutations, the *FoldX* force field remains the most accurate predictor (AUROC = 0.95, Corr = 0.81). Considering two mutations, *Esm_inverse* (AUROC = 0.81, Corr = 0.58) achieves practically equal accuracy of *FoldX* (AUROC = 0.82, Corr = 0.63) and emerges as a highly reliable predictor. For three and four mutations, *Esm_inverse* surpasses other tools, although its performance remains very close to *FoldX* (Fig. 4C and Supp. Fig. 7C). Finally, *Esm_inverse* was confirmed as the best option when considering mutants at all Hamming Distances (AUROC = 0.82) (Supp. Fig. 7B). *TriCombine* performs poorly with few mutations but is very good with 4 mutations category (Fig. 4C, D and Supp. Fig. 7C). It’s worth noting that in both this analysis and the previous on the SH3 domains redesign, *EsmFold* and *Esm_inverse* showed strong correlations with the data. However, *Esm_inverse* tended to overestimate the WT variant, potentially leading to incorrect penalization of variants when using the WT as a reference for identifying more stable designs.

For the Mega-scale dataset, we observed similar results to those seen with the GB1 dataset, with the difference that *Rosetta* is also among the best predictors for point mutations in this case and *FoldX* does not emerge particularly. *ProteinMPNN* (AUROC =0.73, Corr = 0.38) and *Tricombine* (AUROC = 0.77, Corr = 0.38) perform especially well on double mutations in artificial domains, while *FoldX* (AUROC = 0.78, Corr = 0.39) is the best for double mutations in natural domains with crystallographic structures, consistent with the GB1 analysis (Figure 4C, D). In terms of correlation, *FoldX* performs best across all categories, except for natural single mutants, where its performance is essentially equal to *Esm*_*inverse* (Supp. Fig. 7C). Interestingly, performance dropped across all methods when applied to de novo designed proteins and *TriCombine* performs very well with double mutations on artificial domains. However, we should indicate that the data for two mutations should be interpreted cautiously due to the small sample size (Figure 4B). This generalized decline in prediction for de novo designed proteins is most likely due to the use of AlphaFold models as input, which lack crystallographic validation to confirm quality. The lower accuracy is likely because these proteins are artificial and less similar to the natural ones used in AlphaFold’s training datasets.

Looking again at the design goals and their specific questions, which aren’t always tackled directly by general accuracy analysis, the main question is whether, compared to the known properties of the WT, the predictor can forecast one superior variant to be tested experimentally. In this scenario, relying on *FoldX* is the preferred option as, at least for the GB1 dataset, is the only predictor with more true positives than false positives, thus offering a higher probability of obtaining a favorable variant (Fig. 4E). In the case of the Mega-scale analysis none of the methods found more TP than FP (Fig. 4F). This raises the question as to what extent the Mega-scale analysis reflects true changes in stability, or it is heavily contaminated by other factors.

## DISCUSSION

Over the past two decades, computational protein design has struggled to redesign molecules while preserving structural stability, leading many researchers to explore de novo design^40^. While novel molecules offer flexibility, they also raise concerns about immunogenicity and safety, making it advantageous to modify approved proteins for therapeutic use. Effective computational methods that enhance protein properties while maintaining stability would be valuable for advancing protein-based therapies^41^. We selected the most notable and characterized deep learning tools, trained on extensive datasets, and compared them with established force field-based methods and our geometry-based approach.

We assigned our tool *TriCombine* the task of designing the hydrophobic core of a small monomeric domain (SH3), deliberately depriving it of any structural information regarding SH3 domains. We selected the variants between a subset of the top-ranking ones using *FoldX* while ensuring we kept a diversity of predicted structures, ranging from no structural issues to significant Van der Waals clashes. We then used high-quality biochemical stability and structural data at high resolution that we merged with previously determined mutants of the same domain. Most mutants designed over the wild type SH3 structure (1shg) folded well, while those based on the strained multiple mutant 1e6g structure were generally less stable or unfolded. Most mutants designed on the wild type SH3 structure (1shg) folded well, while those based on the strained multiple mutant structure 1e6g were generally less stable or unfolded. As shown in Fig. 1D and 1E, designing mutations on a strained backbone using a rotamer-based approach that cannot adjust the backbone leads to incorrect stability predictions. Notably, mt8, designed over 1e6g to have a cavity, showed significant stability and closed the cavity approximating the WT 1shg structure. Mutant mt18, designed to induce van der Waals clashes, proved more stable than the WT with larger backbone deviations. Remarkably, mutants predicted by *FoldX* without structural issues were the same found to be closer to the WT structure, whereas those intentionally designed with conformational imperfections showed significant changes in the backbone. The fact that SH3 variants are correctly scored by *FoldX* when the structure is modelled by modern AI tools proves the point that the force field works well but accurate modelling of the backbone is required.

When comparing tools’ predictions for the stability of 36 SH3 variants, *EsmFold* and *AlphaFold2* coupled with *FoldX* and *Rosetta* force fields respectively (*EsmFold_F* and *AlphaFold2_R*), demonstrated the highest accuracy in determining variant stability, without significant difference (CI shown in Fig. 2) showing suitability for wild type-based design. *EsmFold* exhibited the strongest correlation, followed by *FoldX* and *AlphaFold2* scores. Both *Esm_inverse* and *ProteinMPNN* struggled to identify variants more stable than the WT, potentially leading to misleading results if solely relied upon. However, when we use *Esm_inverse* to screen the top candidates without reference to the wild type, its correlation shows it can deliver satisfactory results. *ProteinMPNN* showed a weaker correlation, similar to the observations in the following GB1 and Mega-scale analysis. Surprisingly, both *FoldX and Rosetta*, operating with a fixed backbone, achieved higher correlations compared to *RoseTTAFold,* even when coupled with force fields.

this analysis was crucial in demonstrating that *FoldX* and *Rosetta* force fields once again excel at predicting the effects of independent mutations, even when extensively covering mutant proteins (see Supp. Figure 8 and Supp. Table 8). The relatively limited structural data available for hAChE, PTE, and TNfn3 compared to the SH3 domain may explain the performance drop of *EsmFold* and *Esm_inverse*, even when *EsmFold* was combined with force field scorers. The consistent mid-to-high performance of the simple geometrical tool *TriCombine*, which remained relatively unaffected by dataset changes here and in GB1/Mega-scale validations, highlights its potential for further refinement as a versatile inverse folding predictor. Notably, AlphaFold2, when combined with force field functions, proves to be a consistently reliable predictor in all cases.

The structural analysis of 11 high-resolution structures of the designed mutants confirmed *EsmFold* as excellent predictor, at least at modeling the structure of SH3 variants, followed by *AlphaFold2* and *EsmFold* when side chains were relaxed with *FoldX* (*AlphaFold2_r_F* and *EsmFold_r_R*). Methods utilizing *Rosetta* relaxation performed less effectively than modeling side chains over the wild type’s fixed backbone with *FoldX*, once again, possibly due to accurate side chain modeling by *FoldX* at positions like 11, where no backbone adaptations were required. These results aligned with the previous stability predictions, emphasizing the effectiveness of integrating new modeling methods with precise scoring functions.

We also explored a more complex case with greater statistical power: the simultaneous mutation of four interacting residues within the GB1 domain at the interface with a bound IgG protein. In this scenario, we investigated 160,000 mutations and their ability to bind to the IgG target protein, requiring both protein stability and ongoing binding. This imposes an additional constraint compared to the SH3 domain cases discussed above. Here, mutations must not only avoid unfolding the protein but also refrain from significantly disrupting the structure to maintain binding. In this case we find that for single mutations the most accurate method is *FoldX*, while for three and four mutations *EsmFold* is slightly better. If we look at the capability of the predictors to forecast a superior variant than the WT, at least for the GB1 dataset, relying on *FoldX* is again the preferred option, since the higher likelihood of obtaining a favorable variant. However, it’s worth noting that in both this analysis and the one focusing on SH3 domain redesign, *EsmFold* and *Esm_inverse* showed strong correlations with the data. However, *Esm*_*inverse*, like the other inverse folding tool *ProteinMPNN*, tends to overestimate the wild type variant, which can lead to incorrect penalization of desirable alternative designs. Correlation with experimental data helps to assess whether the predictor follows the experimental trend, but when selecting just two or three designs from thousands, what matters most is how reliably the predictor ranks variants relative to a ground truth, such as the wild type.

The Mega-scale dataset broadly confirmed the trends observed in the GB1 analysis, while revealing a performance drop for mutations in artificial domains. This is likely due to the lower quality of the input structures, which were *AlphaFold2* models taken directly from the original publication. As these proteins are artificial, they lack close sequence and structural homologs in the *AlphaFold2* training databases. In contrast, for natural domains, we only used models when they fell within an acceptable RMSD (0.5Å) of an available crystal structure (see Supplementary Note 6). For artificial proteins, no such reference structures were available, limiting model reliability. This helps explain the decline in performance of sequence-agnostic tools like *FoldX* and *Rosetta*, which depend strongly on the accuracy of the input structures. Inverse folding tools perform particularly well on well-represented domains (GB1 or SH3), where structural information from multiple species and rich multiple sequence alignments are available and embedded in their training. Interestingly, *TriCombine* performed better than other tools for double mutations in artificial domains and performed on par with *ProteinMPNN* and *Rosetta* in the four-mutation category of the GB1 dataset, supporting the idea that it was correctly tailored for flexible multi-mutation design. It is worth stressing that every method struggled to separate TPs from FPs in the Mega-scale dataset, suggesting that factors beyond stability mattered here, so these results should be interpreted with caution.

## CONCLUSIONS

In summary, *Rosetta* and *FoldX* force fields show exceptional performance when paired with structural predictors like AlphaFold2 and *EsmFold*, with no significant difference between them. *EsmFold* and *Esm_inverse* proved highly reliable for predicting mutants in well-characterized domains like GB1 and SH3. While they continue to perform well for other domains, their effectiveness becomes comparable to other tools or diminishes in less well-represented domains. *FoldX* remains the best at predicting point mutations. However, when minor conformational changes in the target protein are negligible, a simple geometric superposition model like *TriCombine*, combined with a fundamental scoring function such as *FoldX*, yields promising results, especially when numerous concurrent mutations (>4) are involved, the very situation the method was created for. Further improvements of *TriCombine* could be achieved through implementations of the method in nonlinear scoring functions, as demonstrated in a recent work by Ding et al.^42^ or by extending its application beyond hydrophobic core design to protein-protein interfaces, which will require a dedicated TriX database containing interface triangles defined by clear side chain interactions.

In our investigation, we explored effective combinations of AI modeling tools and their integration with force field functions, highlighting those that enable the modeling community to attain high accuracy.

## MATERIAL AND METHODS

### TriX database generation and querying

The TriXDB was generated starting from a representative set of 5095 high resolution (<1.5Å) crystal structure where entries with more than 95% of similarity were reduced to one representative. Every model was virtually digested in triplets of interacting residues for which geometrical features were registered into a TriX entry of the database. At first, triplet drafts were determined when at least one atom of each amino acid of the triad could find a backbone one from the other two residues within a sphere of 18Å, the largest Cα-Cα distance observed in TriXDB, between two interacting residues. This broad set of triplets was later pruned to those having at least one couple of amino acids with a pair of atoms interacting (less than 4.5Å). When a triplet of residues was determined within a given structure, the set of distances between pairs of α and β carbon of the three residues were registered together with the number of contacts between the three residues and their identities. To stratify the population of TriXs composing the database three levels of contact are assigned: one contact if at least one atom from one residue side chain stays within 4.5Å of any atom of any of the two other residues. Triangles with less than 1 contact were not included in the TriXDB. Two and three contacts follow the same rule, requiring respectively two and three different side chains to contact at least another one within the triplet. We captured 60,563,237 of these natural "building blocks" and incorporated them in the TriXDB precomputing the geometrical relationships. During the saturation process of the *TriCombine* algorithm, features from triangles determined from the input structure are calculated and used to query the TriXDB. Both carbon alpha and beta distances can be searched with a dubiety parameter (respectively “*Cα-dubiety*” and “*Cβ-dubiety*”) allowing deviations from the observed distances, finally a filter based on the number of contacts (“*contacts*”) can be applied to only retrieve triangles with a certain number of internal interactions.

### TriCombine methodology

*TriCombine*’s approach is based on the novel ModelX molecular fragment type TriX. When analyzing a structure for redesign, the distances between input residues and their neighbors (within a distance threshold decided by the user) are measured. Then, all possible triangles are considered (Supplementary Fig. 1A) and the TriXDB is queried with the obtained Cα-Cα and Cβ-Cβ distances (Supplementary Fig.1B,C). Users can exclude TriXs digested from undesired structures and specify the number of contacts that retrieved TriX triangles has to come with, adapting in this way the degree of required interconnectivity between residue positions. Another fundamental querying parameter allows dubiety when looking for compatible Cα-Cα and Cβ-Cβ distances between input structure and database TriXs which considers implicitly possible local backbone moves upon mutation (Supplementary Fig.1B).

When a matching triangle is found in the database, the identities of TriX residues are stored in a substitution matrix (nodeFrequencies), collecting the frequency with which all found amino acids have been proposed by each of the triangles converging on that position (Supplementary Fig.1D). An analogous substitution matrix (edgesFrequencies) stores the frequency of amino acid pairs for each couple of residues forming an edge (Supplementary Fig.1E and Supplementary Note 1). The information provided by the TriX saturation process is then extracted from the substitution matrices to compute two scores for each specific combination of amino acids: edgeScore based on edgeFrequencies and nodeScore based on nodeFrequencies (Supplementary Note 1). In the case of nodeScore, each mutant combination is scored by summing up substitution frequencies for each amino acid at each residue position, as specified by the combination itself. Analogously, the edgeScore equals the sum of the frequencies of amino acid pair substitutions for residues forming an edge (Supplementary Fig. 1F). Through its saturation approximation, *TriCombine* showed to be capable of ranking billions of possible sequences in order to suggest novel stable mutants for a given fold.

*TriCombine* takes in input a PDB structure, a list of residue positions (“*scan-residues*”) and a set of parameters. The tool goes through the input residues and their neighbors collecting them into triplets of residues. Each of those triplets serves as a scanning triangle (triScan) to search for geometrically compatible triXs within the TriXDB database. Each triScan retrieves compatible triXs depending on a set of parameters (Supplementary Note 3), this process that we call “saturation” fills two frequency matrices. The first matrix counts the occurrence of each found amino acidic identity hit at each scan position (node), as in a substitution matrix. The second one collects couples of amino acid substitutions for pairs or positions belonging to the same triScan (edges). To avoid biases, the nodes frequency matrix is normalized by the natural occurrence of amino acids measured in the TriXDB (Supplementary Note 4). All possible scan position sequences are generated, but the user can decide to restrict the considered amino acids to a specific list (with the parameter “*allowed-AAs*”), or to set a saturation threshold per position (with “*combo-th*”). Such threshold allows to limit the list of possible amino acids for a specific position to the most frequent ones (Supplementary Note 3) and reduce the spectrum of sequences to be scored. To avoid biases such frequencies are corrected by natural occurrence of each amino acid as mentioned above. For each generated sequence, node and edge scores are computed respectively as the sum of each amino acid frequency and the sum of each edge frequency. Between the parameters explained in Supplementary Note 3, “*exclude*” allows the user to pass a list of PDBs to exclude: triangles coming from the listed structures won’t be used in the saturation process. The edge score can be calculated in two ways determined by a parameter called “*context*”, that can be set to true or false. When “*context*” is set to false the score calculation for a candidate sequence only considers frequencies of edges between residues listed in input, scoring mutant combinations solely based on the TriXs fitted on the network of scan-residues. When “*context*” is set to true, the contribution of TriXs involving neighbor residues is considered, biasing the score depending on protein context. The saturation process is never affected by this parameter, in all cases the saturation process considers edges between scan-residues and neighbor residues.

### FoldX SH3 candidate modeling

The first 10,000 combinations of mutated proteins were prepared as input files for the *FoldX BuildModel* command. All modeling steps were conducted using a version of the initial protein structure (1shg and 1e6g) firstly energetically minimized with *FoldX RepairPDB* and later mutated to Alanine at the nine specific positions: Val 23, Val 46, Val 9, Ala 11, Val 53, Val 58, Met 25, Leu 31, and Val 44 for 1shg, and Leu 23, Val 46, Val 9, Val 11, Ile 53, Leu 58, Ile 25, Leu 31, and Val 44 for 1e6g. This approach was chosen both to avoid the rotamer positioning being dependent on the WT side chain configuration and to simulate the conditions where contextual residues are to be designed blindly. Each mutant was then evaluated and ranked by *FoldX Stability*. The *FoldX SequenceDetail* tool was used to identify models with less than 2 designed residues having van der Waals clashes above 0.8 kcal/mol, the significance of the *FoldX* force field^43^. These marked combinations were then analyzed with the *FoldX PSSM* algorithm, which explored the potential impact of all 20 single and double mutations on the clashing residues. Finally, the initial 10,000 models, along with their clash-solved variants, were ranked again by *Stability*. The top 500 candidates were then re-modeled in the Alanine version of the starting structures, following 10 different random orders of mutation per candidate. The most stable model from each sequence variant was selected as the representative to reduce the chance of penalizing a sequence variant having its model stuck into an energetically unfavorable conformation. We finally selected 8 variants based on 1shg and 8 based on 1e6g by looking at *FoldX* total energy and *SequenceDetail* per residue energies.

### Protein variants modeling and scoring prediction with predictive tools

In this part of our study, we used 36 unique stability measures for SH3 domain variants, all obtained through chemical denaturation. The data for 18 of these variants was collected from experiments conducted as part of this study, while for the remaining 20 variants was taken from literature^21^. Among the 18 newly produced SH3 proteins, the WT (1shg) and a 5-mutation variant (1e6g), are also part of the older set.

The experimental stabilities were compared with 20 predictions between metrics and combinations:

i. *EsmFold* was installed locally by following the instructions on the GitHub repository (https://github.com/facebookresearch/esm). The ESMFold^44^ model, along with OpenFold^45^, was utilized to generate structures. Only variant sequences were employed to create a single 3D model and their respective scores, corresponding to the average pLDDT, were extracted from the output of the original script provided in the GitHub repository (esmfold.py).
ii. *Esm_inverse*^13^ was coapted from the *Esm* installation at the same github repo (https://github.com/facebookresearch/esm). The script used to generate scores for given sequence variants was feeded with sequences, a 3D structural model of the WT and the chain name to be designed. The original script from the github repo was used (score_log_likelihoods.py).
iii. *ProteinMPNN* (Dauparas et al. 2022) was locally installed from the official repository (https://github.com/dauparas/ProteinMPNN). Variant sequences were scored using the routine provided by the example script (submit_example_3_score_only_from_fasta.sh) from the repository. This involved passing the variant sequence, the 3D model corresponding to the WT, and the molecule name as parameters.
iv. *FoldX* version 5^2^ was employed for various tasks, and all the relevant commands can be found in the official documentation (https://foldxsuite.crg.eu/documentation).
v. *AlphaFold2*^24^ was used through the ColabFold^46^ package (https://github.com/sokrypton/ColabFold) locally installed following the github repo instructions. Structural models for protein variants and their relative scores were obtained launching the “colabfold_batch” command with the variant sequence and the following parameters: “--templates”, “--num-recycle 20”, “--num-models 3”, “--model-type auto”, “--random-seed 16”.
vi. *Rosetta*^1^ was executed using PyRosetta^47^ in three different ways: to generate fixed backbone mutants, evaluate structures with its energy function, and relax structures. Energies were measured using the standard *Rosetta’s* scoring function "Ref2015"^27^. The fixed backbone side chain design was achieved by employing the "FastDesign" mover and setting fixed backbone to the MoveMap, by following the RosettaCommons tutorial https://github.com/RosettaCommons/PyRosetta.notebooks/blob/master/notebooks/06.04-Protein-Design-2.ipynb). ΔΔG values for mutations were calculated by computing the energy difference between the WT model and the mutant one. Relaxation was performed using the "FastRelax" protocol (https://graylab.jhu.edu/PyRosetta.documentation/pyrosetta.rosetta.protocols.relax.html#pyroset ta.rosetta.protocols.relax.FastRelax), guided by the "Ref2015" scoring function, and allowing backbone relaxation.
vii. *FoldX* BuildModel and Stability commands were launched as described in the manual at https://foldxsuite.crg.eu/documentation#manual respectively to generate mutants and evaluate models stabilities.
viii. *RoseTTAFold*^25^ was installed following the instructions provided in the GitHub repository (https://github.com/RosettaCommons/RoseTTAFold), with PyRosetta support. The "run_pyrosetta_ver.sh" script from the original repository was executed, using protein sequences in FASTA format as input.
ix. *Esm_F*, *AlphaFold2_F*, and *RoseTTAFold_F* represent the combination of modeling methods with the *FoldX* force field scoring function. Energy values were obtained using the *FoldX* "Stability" command applied to models generated by the respective methods. Similarly, *Esm_r_F*, *AlphaFold2_r_F*, and *RoseTTAFold_r_F* energy values were obtained with the "Stability" command from *FoldX*, executed after the “RepairPDB” command over the models generated by the respective methods. The same process applies to *Esm_r_R*, *AlphaFold2_r_R*, and *RoseTTAFold_r_R*, but using the “FastRelax” routing from PyRosetta.

A list of all mutants with their denaturation stabilities (kcal/mol) and corresponding predictions is provided in Supplementary Table 5. For the SH3 dataset, two batches were included: the original data from this manuscript and a previously reported dataset from Ventura et al. ^21^. Due to slight discrepancies in ΔG values for the WT variants between the two batches—resulting from different slopes used in the fitting procedure—ΔΔG values were calculated for each mutant relative to its corresponding WT in both batches. The same approach was applied to stability predictions for the scoring methods considered, ensuring to eliminate any bias.

### Sequence alignment of mutants with the whole SH3 Pfam family

To investigate if any of our SH3 domain mutants were already observed we obtained a multiple sequence alignment of the annotated SH3 sequences collected in the Pfam database^36^. The sequence of PDB 1shg was used to determine the family result being PF00018. The “full alignment” for the family was downloaded as well as the full sequences of each alignment entry. The alignment was repeated including 6 and 8 extra residues at N and C terminal respectively to include residues 9 and 58 of 1shg, not represented in Pfam “full alignment”. Using the “seed alignment” sequences of Pfam a multiple sequence alignment was obtained through the software t-coffee^48^ and served as input to Hmmer^49^ for the generation of an HMM model. With *hmmeralign* from Hmmer an alignment of the extended 92,969 SH3 sequences belonging to the family was produced including separately each one of the mutant sequences of the experimental validation of the *TriCombine* algorithm.

### Mutants production and purification

All the ORFs SH3 domain mutants were cloned in a pcoofy2 expression vector^50^, expressed and purified with the following procedure. Plasmid harboring the construct were transformed into *E. coli* Rosetta™(DE3) pLysS Competent Cells (Novagen). Starter cultures were grown in Luria– Bertani (LB) medium, supplemented with 30 μg/ml chloramphenicol or 50 μg/ml kanamycin at 37°C overnight. These cultures were used to inoculate expression cultures in 2TY medium prepared Cell cultures were further cultivated at 37°C and induced at an OD of ∼0.6–0.7 with a final concentration of 0.2 mM IPTG, followed by protein expression at 18°C overnight with subsequent centrifugation to harvest the cells. Pellets were washed with PBS and stored at −80°C or immediately used for protein purification.

Purification was initiated by thawing the pellets on ice with subsequent resuspension in buffer A (20 mM Tris–HCl, 150 mM NaCl, 0.1% (wt/vol) Triton X-100, 1mM DTT, pH 7.4) plus 0.1% (wt/vol) Triton X-100 and cOmplete™ Protease Inhibitor Cocktail (Merk). Cells were lysed with an Emulsiflex high-pressure homogenizer (Avestin) The lysate was clarified by centrifugation and the soluble fraction was added to a HistrapFF 5ml column (Cytiva) (Cytiva) which was previously equilibrated with buffer A. The column was washed stepwise with buffer A supplemented with 5 and 10 mM imidazole. A His-tagged Trx-SH3 fusion protein was eluted with a final elution buffer (20 mM Tris–HCl, 500 mM NaCl, 1mMDTT, 250 mM imidazole, pH 7.4). The elution fractions were pooled and treated overnight with 3 C PreScission Protease in a buffer A at 4C. Subsequently, the sample was loaded again to the Histrap FF column again and the flow thought containing the SH3 protein was dialyzed to the final buffer (PBS1x plus 10% Glycerol and 1mM DTT), concentrated to 5 mg/ml and stored at −80°C until further use.

### Stability assay

All protein solutions in absence or presence of urea (concentrations from 0.5M to 8M) were prepared in buffer 50 mM sodium phosphate pH7.0 using the same protein concentration (9µM). Molar urea concentrations were checked and adjusted by measuring the refractometry index^51^. The mixtures of protein in the mentioned buffer were treated with appropriate denaturant concentration for 1h at 298 K prior to experiment.

Structural changes during protein chemical denaturation have been determined by monitoring intrinsic fluorescence shifts of SH3 tryptophan residues (Trp 41, Trp 42) due to local environment alterations through protein unfolding.

Chemical denaturation experiments of spectrin SH3 and mutants were monitored by tryptophan fluorescence spectroscopy on a Cary Eclipse fluorescence spectrophotometer equipped with a external temperature controller unit at 298 K. Fluorescence emission spectra were recorded in the wavelength range of 305-500 nm with excitation at 290 nm. For all fluorescence measurements, quartz cuvette of 10 mm pathlength was used with excitation and emission slits set at 5 nm band width. Retrieved data have been fitted as explained in Supplementary Note 5, denaturation curves fitted parameters and ΔGs are shown in Supplementary Fig. 2.

### Protein crystallization and X-ray data collection

Ninety-six-well plates crystallization screenings were performed on a Phenix (Art Robbins Instruments) robot mixing 150 nL of reservoir condition with 150 nL of protein solution at the proper concentrations for each mutant (Supplementary Table 6). The optimized crystals of mutants were obtained after mixing 1 µL of each mutant with 1 µL of crystallization condition for each mutant (Supplementary Table 7) in a ratio of 1:1, respectively. Crystals used for data collection were flash frozen in liquid nitrogen with 15% glycerol as cryo-protectant.

X-ray data collection was carried out at Xaloc beamline (ALBA Synchrotron, Spain). Datasets were processed with autoPROC^52^ using XDS^53^, Aimless and Pointless^54^ from the CCP4 suite of programs^55^.

### Determination of crystallographic structures

Crystals belonging to space groups P2_1_2_1_2_1_ and C222_1_ contain one subunit in the asymmetric unit. The structure was solved by molecular replacement with the program Phaser^56^ using as a search model the structure from native SH3 protein (PDB code 1shg). Rebuilding and refinement of all the structures were performed alternating interactive and automatic cycles with programs Coot^57^ and Refmac^58^ from CCP4 and Phenix refine^59^ (Supplementary Table 6). The final refined structures have been deposited in the PDB with codes: 9eyf (mt8), 9ih7 (mt12), 9eyc (mt13), 9eyb (mt15), 9eye (mt16), 9iha (mt17) and 9ih6 (mt18).

### Structure visualization measurements and image rendering

Models and crystals structures were captured using YASARA^60^, also used to calculate models’ cavities, residue volumes and RSMD.

## DATA AVAILABILITY

*TriCombine* is a command-line tool part of the ModelX toolsuite. It’s written in C++ and is designed to be portable across different operating systems, including Linux-64bit, MacOS-64bit, and Windows-32/64bit. The tool makes use of an SQL relational database, which is included in the package as a SQL dump file called PepXDB_5k. PepXDB_5k contains all the structural data used by the software, including the TriX fragments. ModelX is distributed under a proprietary license, it can be downloaded publicly from http://modelx.crg.es. Academic users can get free licenses after registration, commercial licenses are available for for-profit organizations or companies. FoldX is distributed under a proprietary license, it can be downloaded publicly from http://foldx.crg.es. Academic users can get free licenses after registration, commercial licenses are available for for-profit organizations or companies.

## ACKNOWLEDGMENTS

We would like to thank the Centre for Genomic Regulation (CRG) Technology & Business Development Office (TBDO) for support with licensing information, the CRG Tecnologías de Información y Comunicación (TIC) for assistance with web hosting, and the Scientific Information Technologies (SIT) for distributed computing. The Protein Technologies Unit at the Centre for Genomic Regulation. A.M.F.E. acknowledge the Generalitat Valenciana funding Consolidated Research Groups Grant AICO/2020/026. Many thanks are given to the XALOC beamline team at ALBA (Barcelona) for their support during data collection and to the Crystallography Platform at the Barcelona Science Park (PCB). Marc Weber for helping with fitting curves. We appreciate all the feedback from the members of the L.S. lab, especially from Samuel Miravet Verde and Leandro Radusky. This work was supported by funding from the Spanish Ministry of Science and Innovation (Plan Nacional BFU2015-63571-P). We also acknowledge the support of the Spanish Ministry of Science and Innovation to the EMBL partnership, the Centro de Excelencia Severo Ochoa, and the Centres de Recerca de Catalunya (CERCA) Programme / Generalitat de Catalunya. The project that gave rise to these results was supported by a fellowship from “la Caixa’’ Foundation (ID 100010434; fellowship code LCF/BQ/DI19/11730061). D.V thanks a Margarita Salas fellowship.

## AUTHOR CONTRIBUTIONS

D.C, J.D and L.S. conceived the research. D.C and J.D. built TriCombine algorithms and databases. D.C., J.D. and L.S. performed SH3 mutants design. D.C. and L.S. performed the modeling and predictability study of the new SH3 variants. D.C performed the global SH3, GB1 and Mega-scale predictive analysis with the various computer modeling methods. DC and RH performed the analysis on the “bulk” dataset. Protein Technology Unit produced and purified the SH3 mutants. A.M.F.E conducted SH3 stability measurements. D.V. and I.F. solved SH3 mutants structures. D.C., J.D., D.V. and I.F. contributed to the discussion. R.R. curated the software versioning and availability. All authors contributed to writing and editing of the manuscript.

## COMPETING INTEREST

The authors declare no competing interests.

## Notes

### Competing Interest Statement

The authors have declared no competing interest.

## REFERENCES

1. Liu Y, Kuhlman B (2006) RosettaDesign server for protein design. Nucleic Acids Res. 34:W235–8.

2. Delgado J, Radusky LG, Cianferoni D, Serrano L (2019) FoldX 5.0: working with RNA, small molecules and a new graphical interface. Bioinformatics 35:4168–4169.

3. Schymkowitz J, Borg J, Stricher F, Nys R, Rousseau F, Serrano L (2005) The FoldX web server: an online force field. Nucleic Acids Res. 33:W382–8.

4. Pronk S, Páll S, Schulz R, Larsson P, Bjelkmar P, Apostolov R, Shirts MR, Smith JC, Kasson PM, van der Spoel D, et al. (2013) GROMACS 4.5: a high-throughput and highly parallel open source molecular simulation toolkit. Bioinformatics 29:845–854.

5. Zhang Y, Kolinski A, Skolnick J (2003) TOUCHSTONE II: a new approach to ab initio protein structure prediction. Biophys. J. 85:1145–1164.

6. Dantas G, Kuhlman B, Callender D, Wong M, Baker D (2003) A large scale test of computational protein design: folding and stability of nine completely redesigned globular proteins. J. Mol. Biol. 332:449–460.

7. Zheng W, Li Y, Zhang C, Pearce R, Mortuza SM, Zhang Y (2019) Deep-learning contact-map guided protein structure prediction in CASP13. Proteins 87:1149–1164.

8. Wang S, Sun S, Li Z, Zhang R, Xu J (2017) Accurate De Novo Prediction of Protein Contact Map by Ultra-Deep Learning Model. PLoS Comput. Biol. 13:e1005324.

9. Alley EC, Khimulya G, Biswas S, AlQuraishi M, Church GM (2019) Unified rational protein engineering with sequence-based deep representation learning. Nat. Methods 16:1315–1322.

10. Anishchenko I, Pellock SJ, Chidyausiku TM, Ramelot TA, Ovchinnikov S, Hao J, Bafna K, Norn C, Kang A, Bera AK, et al. (2021) De novo protein design by deep network hallucination. Nature 600:547–552.

11. Anand N, Eguchi R, Mathews II, Perez CP, Derry A, Altman RB, Huang P-S (2022) Protein sequence design with a learned potential. Nat. Commun. 13:746.

12. Dauparas J, Anishchenko I, Bennett N, Bai H, Ragotte RJ, Milles LF, Wicky BIM, Courbet A, de Haas RJ, Bethel N, et al. (2022) Robust deep learning-based protein sequence design using ProteinMPNN. Science 378:49–56.

13. Hsu C, Verkuil R, Liu J, Lin Z, Hie B, Sercu T, Lerer A, Rives A (2022) Learning inverse folding from millions of predicted structures. bioRxiv [Internet]. Available from: http://biorxiv.org/lookup/doi/10.1101/2022.04.10.487779

14. Lin Z, Akin H, Rao R, Hie B, Zhu Z, Lu W, Smetanin N, Verkuil R, Kabeli O, Shmueli Y, et al. (2023) Evolutionary-scale prediction of atomic-level protein structure with a language model. Science 379:1123–1130.

15. Watson JL, Juergens D, Bennett NR, Trippe BL, Yim J, Eisenach HE, Ahern W, Borst AJ, Ragotte RJ, Milles LF, et al. (2023) De novo design of protein structure and function with RFdiffusion. Nature 620:1089–1100.

16. Singh A (2024) Chroma is a generative model for protein design. Nat. Methods 21:10.

17. Lin Z, Akin H, Rao R, Hie B, Zhu Z, Lu W, Smetanin N, Verkuil R, Kabeli O, Shmueli Y, et al. (2023) Evolutionary-scale prediction of atomic-level protein structure with a language model. Science 379:1123–1130.

18. Notin P, Kollasch AW, Ritter D, van Niekerk L, Paul S, Spinner H, Rollins N, Shaw A, Weitzman R, Frazer J, et al. (2023) ProteinGym: Large-Scale Benchmarks for Protein Design and Fitness Prediction. bioRxiv [Internet]. Available from: 10.1101/2023.12.07.570727

19. Wu NC, Dai L, Olson CA, Lloyd-Smith JO, Sun R (2016) Adaptation in protein fitness landscapes is facilitated by indirect paths. Elife [Internet] 5. Available from: 10.7554/eLife.16965

20. Tsuboyama K, Dauparas J, Chen J, Laine E, Mohseni Behbahani Y, Weinstein JJ, Mangan NM, Ovchinnikov S, Rocklin GJ (2023) Mega-scale experimental analysis of protein folding stability in biology and design. Nature 620:434–444.

21. Ventura S, Vega MC, Lacroix E, Angrand I, Spagnolo L, Serrano L (2002) Conformational strain in the hydrophobic core and its implications for protein folding and design. Nat. Struct. Biol. 9:485–493.

22. Musacchio A, Noble M, Pauptit R, Wierenga R, Saraste M (1992) Crystal structure of a Src-homology 3 (SH3) domain. Nature 359:851–855.

23. Baek M, DiMaio F, Anishchenko I, Dauparas J, Ovchinnikov S, Lee GR, Wang J, Cong Q, Kinch LN, Schaeffer RD, et al. (2021) Accurate prediction of protein structures and interactions using a three-track neural network. Science 373:871–876.

24. Jumper J, Evans R, Pritzel A, Green T, Figurnov M, Ronneberger O, Tunyasuvunakool K, Bates R, Žídek A, Potapenko A, et al. (2021) Highly accurate protein structure prediction with AlphaFold. Nature 596:583–589.

25. Baek M, DiMaio F, Anishchenko I, Dauparas J, Ovchinnikov S, Lee GR, Wang J, Cong Q, Kinch LN, Schaeffer RD, et al. (2021) Accurate prediction of protein structures and interactions using a three-track neural network. Science 373:871–876.

26. Shanker VR, Bruun TUJ, Hie BL, Kim PS (2023) Inverse folding of protein complexes with a structure-informed language model enables unsupervised antibody evolution. bioRxiv [Internet]. Available from: 10.1101/2023.12.19.572475

27. Alford RF, Leaver-Fay A, Jeliazkov JR, O’Meara MJ, DiMaio FP, Park H, Shapovalov MV, Renfrew PD, Mulligan VK, Kappel K, et al. (2017) The Rosetta All-Atom Energy Function for Macromolecular Modeling and Design. J. Chem. Theory Comput. 13:3031–3048.

28. Goldenzweig A, Goldsmith M, Hill SE, Gertman O, Laurino P, Ashani Y, Dym O, Unger T, Albeck S, Prilusky J, et al. (2016) Automated Structure- and Sequence-Based Design of Proteins for High Bacterial Expression and Stability. Mol. Cell 63:337–346.

29. Kim DN, Jacobs TM, Kuhlman B (2016) Boosting protein stability with the computational design of β-sheet surfaces. Protein Sci. 25:702–710.

30. Dauparas J, Anishchenko I, Bennett N, Bai H, Ragotte RJ, Milles LF, Wicky BIM, Courbet A, de Haas RJ, Bethel N, et al. (2022) Robust deep learning-based protein sequence design using ProteinMPNN. Science 378:49–56.

31. Liu Y, Kuhlman B (2006) RosettaDesign server for protein design. Nucleic Acids Res. 34:W235–8.

32. Blanco JD, Radusky L, Climente-González H, Serrano L (2018) FoldX accurate structural protein-DNA binding prediction using PADA1 (Protein Assisted DNA Assembly 1). Nucleic Acids Res. 46:3852–3863.

33. Blanco JD, Radusky LG, Cianferoni D, Serrano L (2019) Protein-assisted RNA fragment docking (RnaX) for modeling RNA–protein interactions using ModelX. Proceedings of the National Academy of Sciences [Internet] 116:24568–24573. Available from: 10.1073/pnas.1910999116

34. Cianferoni D, Radusky LG, Head SA, Serrano L, Delgado J (2020) ProteinFishing: a protein complex generator within the ModelX toolsuite. Bioinformatics 36:4208–4210.

35. Berman HM, Westbrook J, Feng Z, Gilliland G, Bhat TN, Weissig H, Shindyalov IN, Bourne PE (2000) The Protein Data Bank. Nucleic Acids Res. 28:235–242.

36. Mistry J, Chuguransky S, Williams L, Qureshi M, Salazar GA, Sonnhammer ELL, Tosatto SCE, Paladin L, Raj S, Richardson LJ, et al. (2021) Pfam: The protein families database in 2021. Nucleic Acids Res. 49:D412–D419.

37. Delgado J, Reche R, Cianferoni D, Orlando G, van der Kant R, Rousseau F, Schymkowitz J, Serrano L (2025) FoldX force field revisited, an improved version. Bioinformatics [Internet] 41. Available from: 10.1093/bioinformatics/btaf064

38. Mower JP (2005) PREP-Mt: predictive RNA editor for plant mitochondrial genes. BMC Bioinformatics 6:96.

39. Mower JP (2005) PREP-Mt: predictive RNA editor for plant mitochondrial genes. BMC Bioinformatics 6:96.

40. Huang P-S, Boyken SE, Baker D (2016) The coming of age of de novo protein design. Nature 537:320–327.

41. Listov D, Goverde CA, Correia BE, Fleishman SJ (2024) Opportunities and challenges in design and optimization of protein function. Nat. Rev. Mol. Cell Biol. [Internet]. Available from: 10.1038/s41580-024-00718-y

42. Ding D, Shaw AY, Sinai S, Rollins N, Prywes N, Savage DF, Laub MT, Marks DS (2024) Protein design using structure-based residue preferences. Nat. Commun. 15:1639.

43. Guerois R, Nielsen JE, Serrano L (2002) Predicting changes in the stability of proteins and protein complexes: a study of more than 1000 mutations. J. Mol. Biol. 320:369–387.

44. Lin Z, Akin H, Rao R, Hie B, Zhu Z, Lu W, Smetanin N, Verkuil R, Kabeli O, Shmueli Y, et al. (2022) Evolutionary-scale prediction of atomic level protein structure with a language model. bioRxiv [Internet]. Available from: http://biorxiv.org/lookup/doi/10.1101/2022.07.20.500902

45. Ahdritz G, Bouatta N, Kadyan S, Xia Q, Gerecke W, O’Donnell TJ, Berenberg D, Fisk I, Zanichelli N, Zhang B, et al. (2022) OpenFold: Retraining AlphaFold2 yields new insights into its learning mechanisms and capacity for generalization. bioRxiv [Internet]. Available from: http://biorxiv.org/lookup/doi/10.1101/2022.11.20.517210

46. Mirdita M, Schütze K, Moriwaki Y, Heo L, Ovchinnikov S, Steinegger M (2022) ColabFold: making protein folding accessible to all. Nat. Methods 19:679–682.

47. Chaudhury S, Lyskov S, Gray JJ (2010) PyRosetta: a script-based interface for implementing molecular modeling algorithms using Rosetta. Bioinformatics 26:689–691.

48. Garriga E, Di Tommaso P, Magis C, Erb I, Mansouri L, Baltzis A, Floden E, Notredame C (2021) Multiple Sequence Alignment Computation Using the T-Coffee Regressive Algorithm Implementation. Methods Mol. Biol. 2231:89–97.

49. Eddy SR (2011) Accelerated Profile HMM Searches. PLoS Computational Biology [Internet] 7:e1002195. Available from: 10.1371/journal.pcbi.1002195

50. Scholz J, Besir H, Strasser C, Suppmann S (2013) A new method to customize protein expression vectors for fast, efficient and background free parallel cloning. BMC Biotechnol. 13:12.

51. Warren JR, Gordon JA (1966) On the refractive indices of aqueous solutions of urea. J. Phys. Chem. 70:297–300.

52. Vonrhein C, Flensburg C, Keller P, Sharff A, Smart O, Paciorek W, Womack T, Bricogne G (2011) Data processing and analysis with the autoPROC toolbox. Acta Crystallogr. D Biol. Crystallogr. 67:293–302.

53. Kabsch W (2010) XDS. Acta Crystallogr. D Biol. Crystallogr. 66:125–132.

54. Evans P (2006) Scaling and assessment of data quality. Acta Crystallogr. D Biol. Crystallogr. 62:72–82.

55. Winn MD, Ballard CC, Cowtan KD, Dodson EJ, Emsley P, Evans PR, Keegan RM, Krissinel EB, Leslie AGW, McCoy A, et al. (2011) Overview of the CCP4 suite and current developments. Acta Crystallogr. D Biol. Crystallogr. 67:235–242.

56. Cao L, Coventry B, Goreshnik I, Huang B, Sheffler W, Park JS, Jude KM, Marković I, Kadam RU, Verschueren KHG, et al. (2022) Design of protein-binding proteins from the target structure alone. Nature 605:551–560.

57. Emsley P, Cowtan K (2004) Coot: model-building tools for molecular graphics. Acta Crystallogr. D Biol. Crystallogr. 60:2126–2132.

58. Murshudov GN, Vagin AA, Dodson EJ (1997) Refinement of macromolecular structures by the maximum-likelihood method. Acta Crystallogr. D Biol. Crystallogr. 53:240–255.

59. Afonine PV, Poon BK, Read RJ, Sobolev OV, Terwilliger TC, Urzhumtsev A, Adams PD (2018) Real-space refinement in PHENIX for cryo-EM and crystallography. Acta Crystallogr D Struct Biol 74:531–544.

60. Krieger E, Koraimann G, Vriend G (2002) Increasing the precision of comparative models with YASARA NOVA--a self-parameterizing force field. Proteins 47:393–402.

61. Therapeutic monoclonal antibodies approved or in review in the EU or US.

62. www.antibodysociety.org/resources/approved-antibodies.

