## supplementary material for "Artificial Intelligence And First Principle Methods In Protein Redesign: A Marriage Of Convenience?"

### Supplementary information

Supplementary Note 1
Supplementary Note 2
Supplementary Note 3
Supplementary Note 4
Supplementary Note 5
Supplementary Note 6
Supplementary Figures 1 to 7
Supplementary Tables 1 to 8

Executable, databases and instruction files for testing of tricombine are available at:
https://www.dropbox.com/scl/fo/dfgaxpbqhv792vvplu3m0/APPfVBrpm8rFr7P0MaJMvcA?rlkey=aq2savm0nbvezjic4c3v1isdy&dl=0

#### Supplementary Note 1

**_TriCombine_ output for 1shg and 1e6g: edgeScores and nodeScores**
EdgeScore and nodeScore are the two output files for TriCombine scores. A combination of residues is scored summing up the frequency for edges connecting two connected substitutions (edgeScore) or single amino acid substitutions (nodeScore). See example in Supp. Fig. 1F.

#### Supplementary Note 2

**SH3 mutants alignment with SH3 domain family**
The sequence of src-homology 3 (SH3) domain was extracted directly from the PDB file 1SHG (KELVLALYDYQEKSPREVTMKKGDILTLLNSTNKDWWKVEVNDRQGFVPAAYVKKLD).
The sequence was later prompted as a search term in Pfam (http://pfam.xfam.org/)(1), showing that it belongs to the SH3_1 family (PF00018). The Pfam "full alignment" of the PF00018 family was downloaded. All sequences in the alignment were extended by 6 and 8 residues respectively at N and C termini. This manipulation was necessary since the original Pfam alignment did not include residues corresponding to 9 and 58 of 1SHG structure, which is part of the domain's hydrophobic core. We proceeded by emulating the Pfam family alignment generation. We aligned 55 extended seed sequences using T-COFFEE multiple sequence alignment (MSA) algorithm(2); with this alignment, a Hidden Markov Model (HMM) was built using *hmmbuild* from HMMER 3.2.1(3). A new alignment was generated for each of the 16 designed mutants by using HMMER 3.2.1 *hmmalign* algorithm and the produced HMM model. Each alignment contained the 92,969 extended sequences belonging to PF00018 and one of the 16 candidates.
Subsequently, the set of alignments was analyzed to gather data about the frequencies of amino acids at the nine hydrophobic core positions that underwent re-design. The highest number of core amino acid identities shared between a mutant and any element in the SH3_1 family MSA was recorded, along with the count of this value (Supplementary Table 2). This information

indicates the largest subset of core amino acids that are common to at least one family member and how many times they occur. None of the designed mutants had an identical core combination with any element of the family alignment.

### Supplementary Note 3

##### *TriCombine* parameters

The *TriCombine* algorithm has various parameters influencing both the analysis results and output format. TriScan objects are triangles of residues used to gather geometrically compatible TriXs from the TriXDB. Compatibility parameters are used to restrict or widen TriXs retrieval.
Briefly, the available parameters with their standard values are:

1. Scan-residues: the comma-separated list of residue positions to be explored by the algorithm in the following format: one letter amino acid - molecule name - residue number [example: AA1,CA2,DB1]
2. CA-dubiety: the allowed dubiety of carbon alpha distance for database's triangles to be matched with the input structure ones.
3. CB-dubiety: the allowed dubiety of carbon beta distance for database's triangles to be matched with the input structure ones.
4. Contacts-th: the minimum number of contacts within TriXs retrieved from the database. 0 means any number is accepted, 1 means at least one residue is contacting another within the retrieved TriX, 2 at least 2 residues are contacting one residue of the TriX. 3 means all residues belonging to the TriX are contacting another residue part of the TriX.
5. Combo-th: this threshold goes from 0 to 1 and represents the degree of saturation satisfaction for each residue position. When equal to one each of the 20 amino acids will be taken into consideration when generating the scored combinations of substitutions. When set to 0.5, amino acid substitutions accepted for each residue position will be limited to the most frequent ones until the total frequency of saturation satisfies the 50% of the amino acid observations for that position.
6. Neigh-th: this threshold is the max distance between residue atoms belonging to the input structure that is considered to make a triScan. If it is set to 8, all residues having an atom in a sphere of 8Å around each scan-residue position are considered to be part of a triScan.
7. allowed-AAs: a string containing single letter amino acids that are allowed to be found in the TriX search during the saturation. If not specified all 20 amino acids are allowed, if by example only "A" is specified then the only TriXs to be retrieved will be those composed by triplets of Ala. If only "AC" are specified, the only allowed TriXs will be those containing either A or C or both at each of the three positions composing a TriX.
8. Context: when True, this parameter makes the computed edge score for mutant combinations to depend on the saturation of triScans containing residues not listed between the scan-residues with the condition that at least one of the residues of the triScan is listed. This parameter makes the score dependent on the structural context that the user does not want to modify. When set to false, the score only depends on the strength of determined scan-residues connections.
9. Frq-only: When equal to True, the scores are not computed for any combination, and only the saturation matrices are produced.

10. Print-Trix: an output file is printed containing all TriX ID information from the TriXDB that hits each scan residue; it also indicates the amino acid of the TriX.
11. Exclude: this optional parameter can be assigned with a filename containing a list of PDB identifiers to be excluded during the saturation process. Any TriX belonging to a PDB file listed here won't be used for the saturation.

### Supplementary Note 4

**Amino acid frequency correction**

When calculating amino acid frequency for position after the triangle saturation process, each amino acid contribution is weighted based on the absolute frequency of such amino acid in the TriXDB:

| | | | | |
|---|---|---|---|---|
| A = 0.08757 | H = 0.02376 | T = 0.05697 | C = 0.01278 | M = 0.01864 |
| G = 0.07969 | S = 0.05950 | D = 0.05991 | K = 0.05595 | V = 0.07214 |
| P = 0.04677 | N = 0.04457 | L = 0.08720 | Y = 0.03595 | E = 0.06253 |
| R = 0.04789 | I = 0.05573 | W = 0.01524 | Q = 0.03731 | F = 0.03979 |

### Supplementary Note 5

**Denaturation curves for SH3 mutants**

Denaturation curves for the 16 proteins designed in this work are shown in Supplementary Figure 1, together with the ones for the WT 1SHG and the expanded variant "1E6G". Curve points are the ratio between fluorescence emission (see Methods) at 335 nm and 400 nm.

Ratio values together with urea concentrations were used to fit the curves, using the following equation from Viguera et al. 1994(4):

$$F = \frac{(F_N + a[U]) + (F_U + b[U]) \times e^{\frac{m[U] - \Delta G_{H2O}}{RT}}}{1 + e^{\frac{m[U] - \Delta G_{H2O}}{RT}}}$$

"F" is the fluorescence value at a certain concentration of urea ("U"), while "$F_N$" and "$F_U$" are the fluorescence values for the fully folded and unfolded states, respectively, in the absence of urea. The free energy of unfolding proteins in the presence of urea is linearly related to the urea concentration(5). $G_{H2O}$ is the apparent free energy of unfolding in the absence of urea. "m" is the proportionality constant that reflects the cooperativity of the transition.

Fitting was performed using Python's scipy library, which uses non-linear least squares. Standard deviation of estimated parameters "m3" (slope at inflection) and "m4" (x axis value at inflection point) are given, as well as "m3" and "m4" covariance. As the stability (ΔG) equals the products of “m3” and “m4”, we obtain ΔG errors as the square root of ΔG variance with the following formula that requires “m3” and “m4” standard deviations (σ) and “m3” and “m4” covariances:

$$\sigma \approx |f| \sqrt{(\frac{\sigma_{m3}}{m3})^2 + (\frac{\sigma_{m4}}{m4})^2 + 2\frac{\sigma_{m3\,m4}}{m3 \times m4}}$$

### Supplementary Note 6

**Preparation of filtered mega-scale datasets**

We extracted 325,132 ΔG measurements from 17,093 sites across 365 domains from the original publication (“Dataset3”)(6) focusing on high-quality data with available ΔΔG estimates. Dataset3 already flagged unreliable ΔΔG values with a “-”, and these variants, along with double mutants not marked with a “pair_name” (indicating non-interacting residues), were excluded. Duplicated wild-type entries were grouped for simplicity and point mutation values obtained over other point mutation variants were excluded due to the lack of available structures as well as insertion and deletion variants. The “WT_cluster” column was used to distinguish natural proteins from artificial ones. Natural proteins in this subset were searched for available high-resolution crystallographic structures (resolution < 2.5Å). Proteins without structures were excluded. The obtained structures were compared with AlphaFold models from the original publication, and models with an RMSD < 0.5Å (after structural alignment using MUSTANG (7)) were retained to ensure reliable predictions. All artificial variant data were retained, and AlphaFold models were used without further filtering. The final dataset contained 163,555 entries from 146 artificial domains and 33 natural ones.

### Supplementary figures

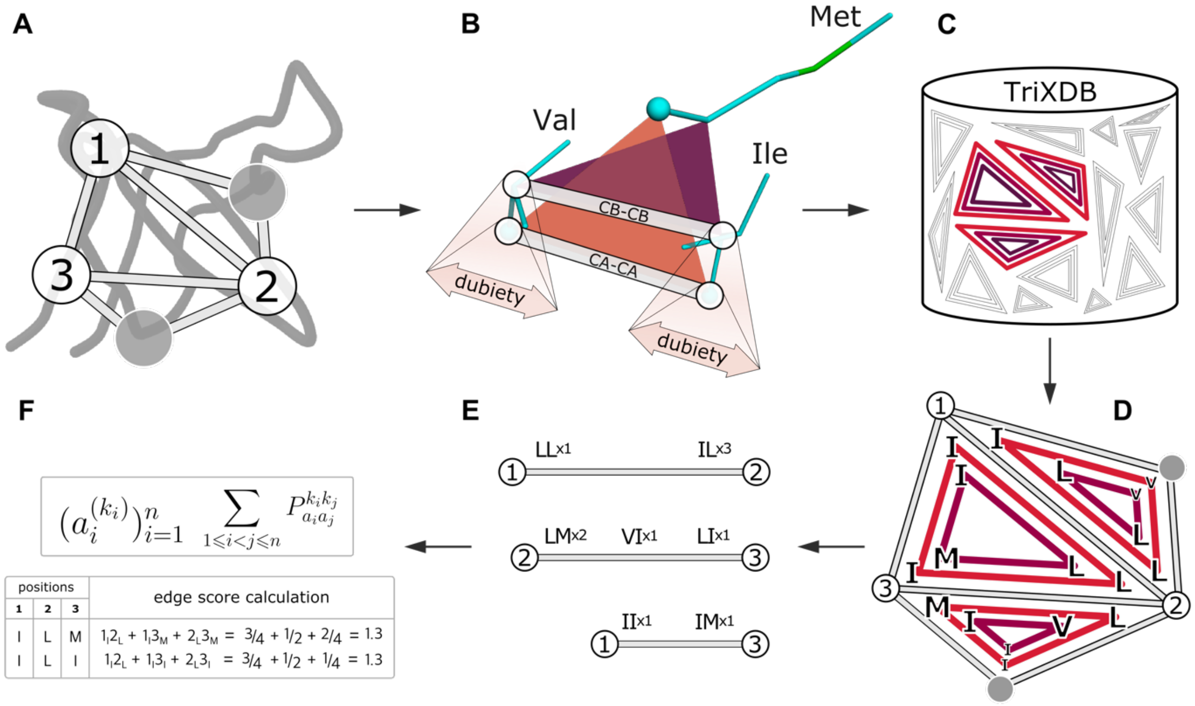


**Supplementary Figure. 1: TriCombine methodology:** A) a protein backbone structure of input and a set of residue positions to be explored (1-2-3) are used, together with neighbor residues (gray circles), to derive all possible triangles geometries. B) TriXs from the TriXDB are compared with query triangles using Cα and Cβ distances, allowing a dubiety. C-D) Many different TriXs can be found with similar geometries and bringing various combinations of amino acid identities (red shades triangles). E) The saturation process stores frequencies of substitution pairs. As an example, the edge between position 1 and 2 is covered by four different TriXs which bring respectively three “IL” substitution pairs and one “LL” pair. F)
ai: residue position; ki: amino acid; P: edgeScore of a combination. For a given mutant combination at positions 1,2,3, its edgeScore is computed as the sum of the registered frequencies of the resulting amino acid pairs. For the combination “ILM” at positions 1,2,3, the edgeScore is the sum of “IL” frequency at 1-2 + “IM” frequency at 1-3 +”LM” frequency at 2-3.

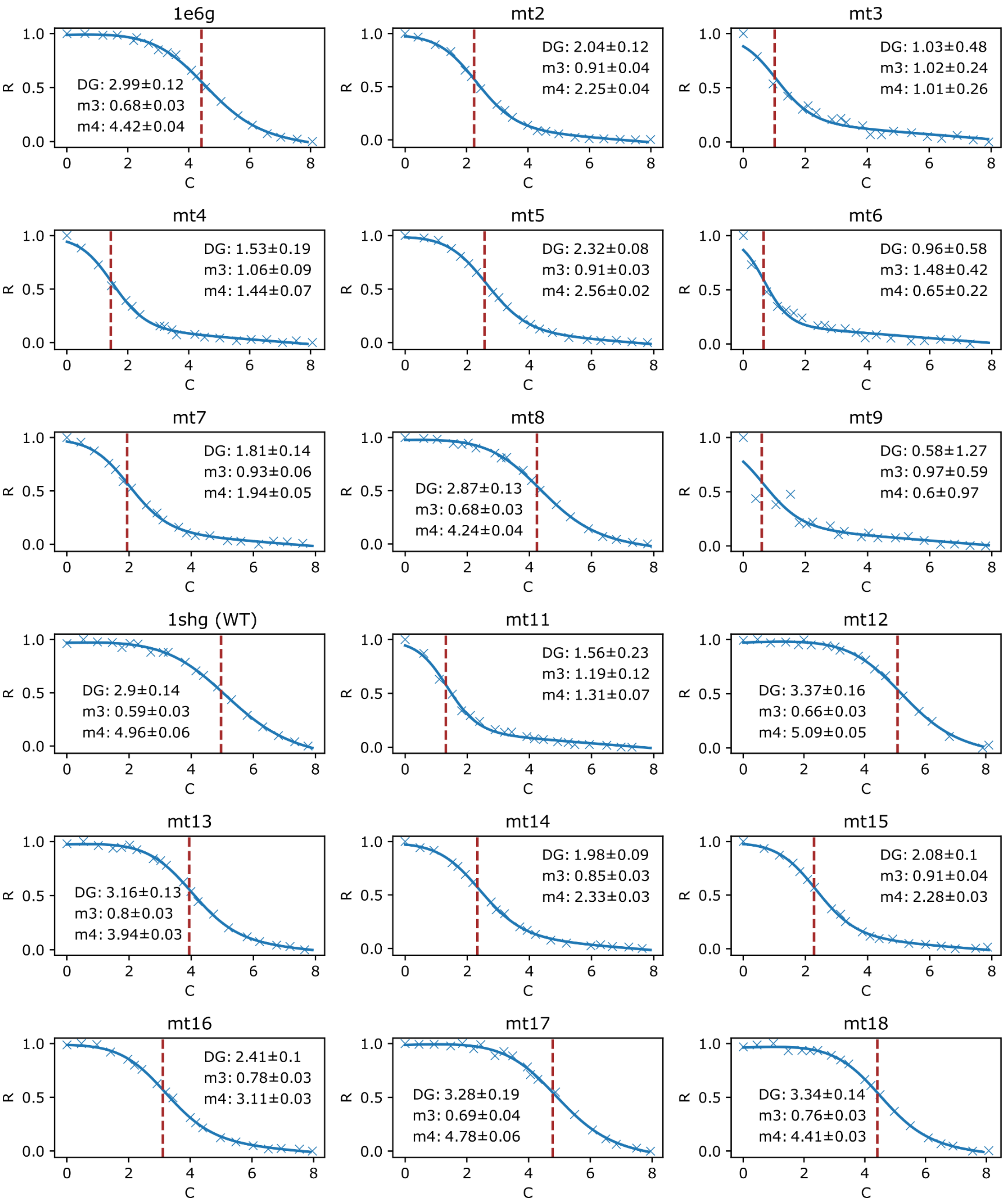


**Supplementary Figure 2: Urea denaturation curves**

Fitting curves for the 16 mutants generated in this work, the WT and “1E6G” variants. The X axis is showing urea concentration (C), the Y axis the ratio between fluorescence emission (R) normalized, a dotted red line shows the fitted m4 value. Each plot contains fitting values for the denaturation curves including the computed ∆Gs in kcal/mol.

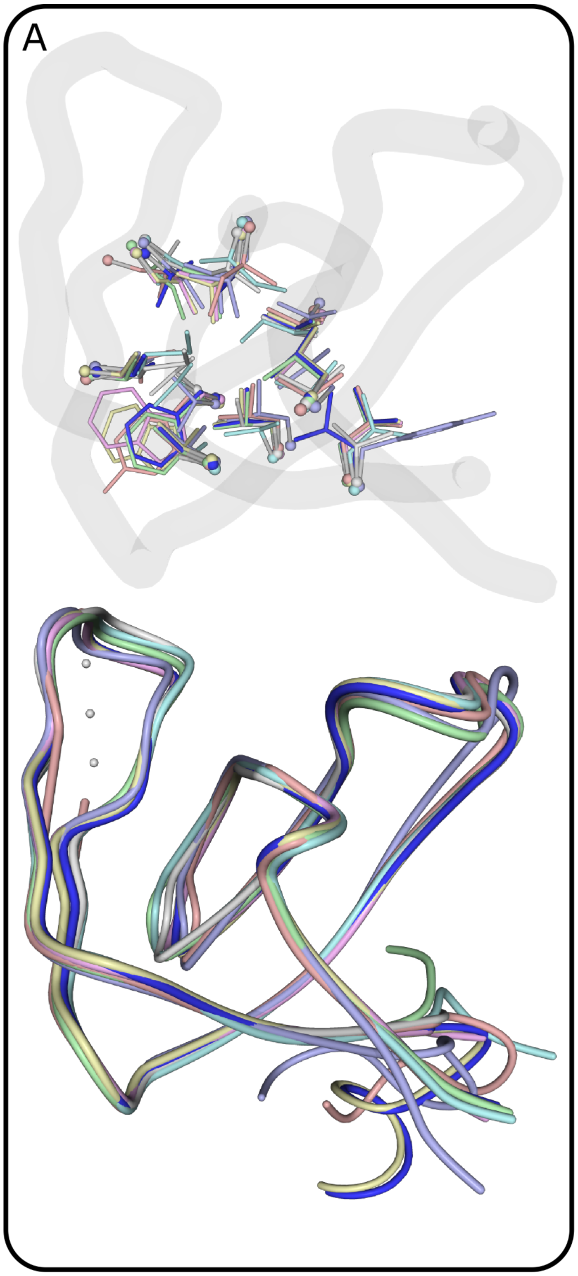


B

| pos | mt8 | | mt12 | mt13 | mt15 | mt16 | mt17 | mt18 |
|---|---|---|---|---|---|---|---|---|
| | 1shg | 1e6g | | | | | | |
| wt | 0.7 | 0.9 | 0.7 | 0.7 | 0.7 | 0.4 | 0.6 | 1.0 |

C

| pos | mt8 | | mt12 | mt13 | mt15 | mt16 | mt17 | mt18 |
|---|---|---|---|---|---|---|---|---|
| | 1shg | 1e6g | | | | | | |
| 9 | 0.4 | 0.3 | 0.5 | 0.3 | 0.3 | 0.3 | 0.4 | 0.3 |
| 11 | 0.2 | 0.3 | 0.2 | 0.2 | 0.3 | 0.2 | 0.3 | 0.1 |
| 23 | 1.9 | 2.1 | 0.6 | 0.8 | 0.9 | 0.0 | 0.7 | 0.6 |
| 25 | 0.8 | 1.1 | 0.6 | 0.9 | 0.9 | 0.1 | 0.5 | 0.5 |
| 31 | 0.3 | 1.1 | 0.2 | 0.2 | 0.2 | 0.3 | 0.1 | 0.6 |
| 44 | 0.4 | 0.5 | 0.3 | 0.5 | 0.4 | 0.2 | 0.3 | 0.7 |
| 46 | 1.1 | 1.3 | 1.4 | 1.6 | 1.8 | 0.3 | 1.3 | 1.1 |
| 53 | 0.6 | 0.6 | 0.4 | 0.3 | 0.2 | 0.5 | 0.3 | 0.6 |
| 58 | 0.4 | 0.8 | 0.2 | 0.3 | 0.3 | 0.2 | 0.2 | 0.3 |
| all | 0.8 | 1.0 | 0.6 | 0.7 | 0.8 | 0.3 | 0.6 | 0.6 |

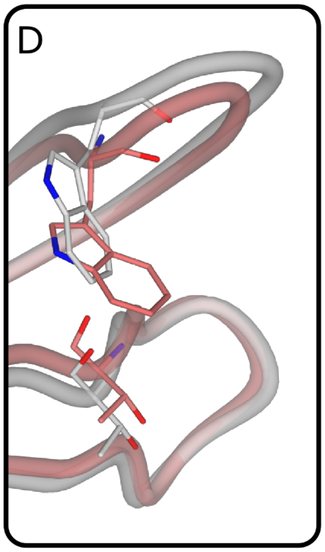


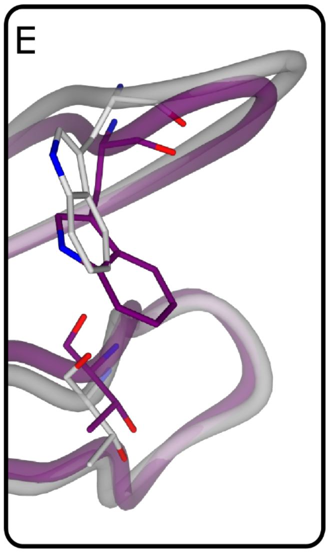


**Supplementary Figure 3. SH3 mutant structures comparisons and RMSDs:** A) superposition of the 7 SH3 mutant structures, showing side chains (top) and backbones (bottom) variability. B) All residue Cα RMSDs of the 7 crystallized mutants with respect to their relative reference structures (1shg and 1e6g). C) Cα RMSDs of SH3 hydrophobic core residues for the 7 crystallized mutants with respect to their relative reference structure (1shg and 1e6g). D) Cα RMSDs of SH3 hydrophobic core residues for the 7 crystallized mutants with respect to their relative AF2 models. E) The superposition between 1shg (gray), mt13 (purple) and mt15 (red) crystals reveals a backbone displacement at position 46 for both mutants, result of the desolvation on the CO group at residue 24 induced by the Trp mutation at position 46.

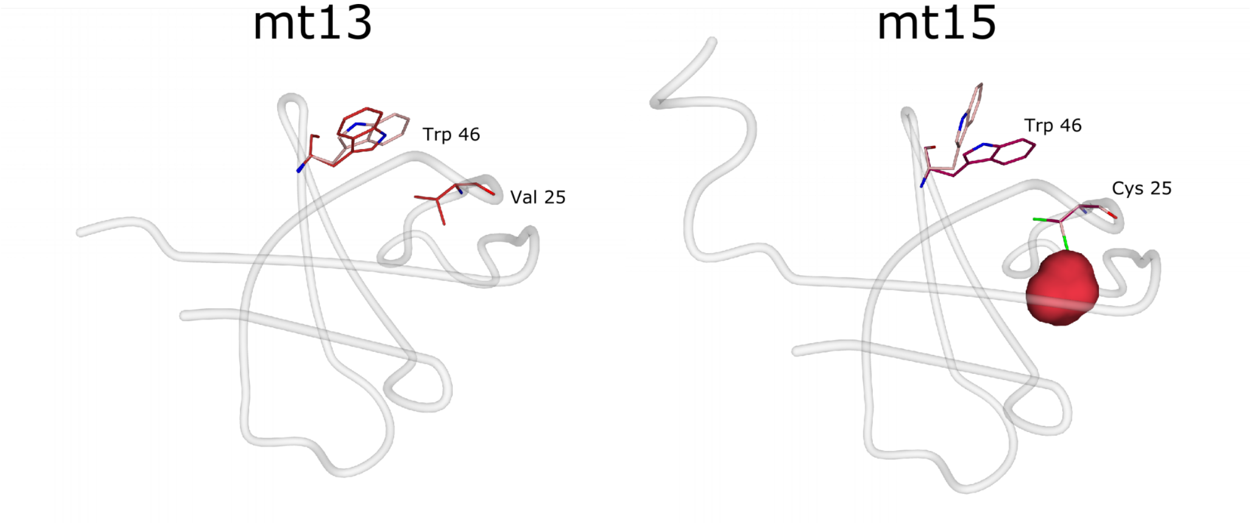


**Supplementary Figure 4: Effect of Tryptophan 46 substitution in mt13 and mt15 crystals**
Details of mt13 and mt15 conformational changes are shown.

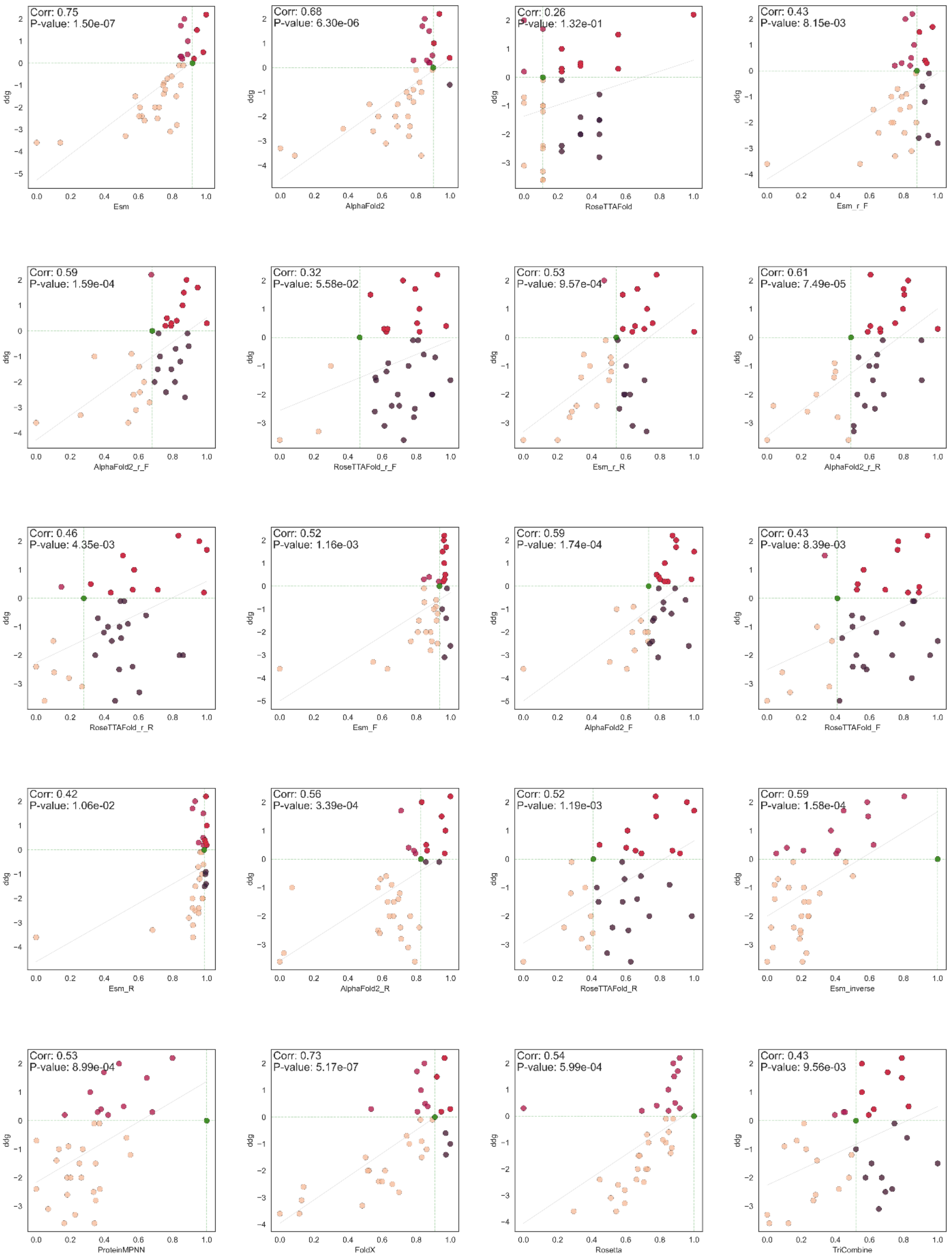
Corr: 0.75
P-value: 1.50e-07
Esm
ddg
Corr: 0.68
P-value: 6.30e-06
AlphaFold2
ddg
Corr: 0.26
P-value: 1.32e-01
RoseTTAFold
ddg
Corr: 0.43
P-value: 8.15e-03
Esm_r_F
ddg
Corr: 0.59
P-value: 1.59e-04
AlphaFold2_r_F
ddg
Corr: 0.32
P-value: 5.58e-02
RoseTTAFold_r_F
ddg
Corr: 0.53
P-value: 9.57e-04
Esm_r_R
ddg
Corr: 0.61
P-value: 7.49e-05
AlphaFold2_r_R
ddg
Corr: 0.46
P-value: 4.35e-03
RoseTTAFold_r_R
ddg
Corr: 0.52
P-value: 1.16e-03
Esm_F
ddg
Corr: 0.59
P-value: 1.74e-04
AlphaFold2_F
ddg
Corr: 0.43
P-value: 8.39e-03
RoseTTAFold_F
ddg
Corr: 0.42
P-value: 1.06e-02
Esm_R
ddg
Corr: 0.56
P-value: 3.39e-04
AlphaFold2_R
ddg
Corr: 0.52
P-value: 1.19e-03
RoseTTAFold_R
ddg
Corr: 0.59
P-value: 1.58e-04
Esm_inverse
ddg
Corr: 0.53
P-value: 8.99e-04
ProteinMPNN
ddg
Corr: 0.73
P-value: 5.17e-07
FoldX
ddg
Corr: 0.54
P-value: 5.99e-04
Rosetta
ddg
Corr: 0.43
P-value: 9.56e-03
TriCombine
ddg

**Supplementary Figure 5: Scatter plots and correlations between unfolding stability for 36 SH3 mutants VS predictions**

The figure shows all scatterplots for 36 SH3 domain mutant stabilities measured experimentally through urea denaturation against predictions performed with the 20 between scoring functions and their combinations (TriCombine, Esm-inverse, proteinMPNN, FoldX, Rosetta, Esm, RoseTTAFold, AlphaFold2, Esm_F, RoseTTAFold_F, AlphaFold2_F, Esm_R, RoseTTAFold_R, AlphaFold2_R, Esm_r_F, RoseTTAFold_r_F, AlphaFold2_r_F, Esm_r_R, RoseTTAFold_r_R, AlphaFold2_r_R). The scatterplots display the experimentally measured ΔΔG values for each variant, calculated by subtracting the wild-type values (1shg and mt10) from the mutant one in the two experimental batches (Ventura and Cianferoni), each with slightly different wild-type values. These are compared to the predicted Δ-scores, computed by subtracting the predicted wild-type value from the prediction for each mutant. The prediction values have been normalized for improved visualization given the variagate set of scales they span through.

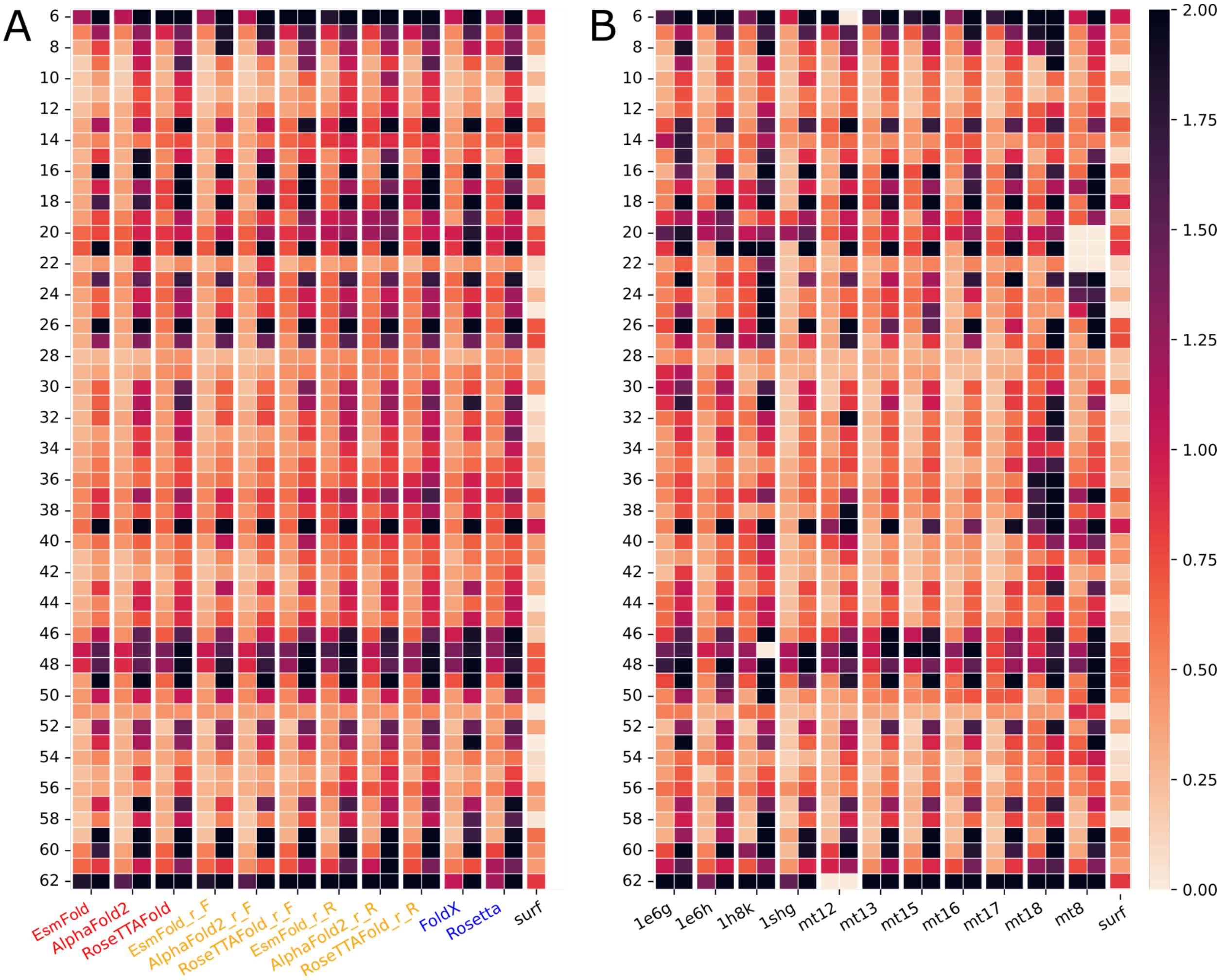


**Supplementary Figure 6: RMSD heatmap Heatmap for SH3 variants averaged over the different methods tested and over the crystallographic structures obtained**

Supp. Figure 6A displays a heatmap, capped at 2Å of RMSD, averaged across the 11 available crystallographic structures of SH3 mutants. Each method is represented by two boxes: the left box shows the RMSD for Cα atoms, while the right one indicates the RMSD for all side chain atoms. These visualizations illustrate how well, on average, each method predicted the position of each residue, thereby highlighting which method performed better for specific positions. In Figure 6B, the average RMSDs for all methods are depicted for each of the 11 structures. This presentation demonstrates the overall predictability of all positions within the domain for each variant.

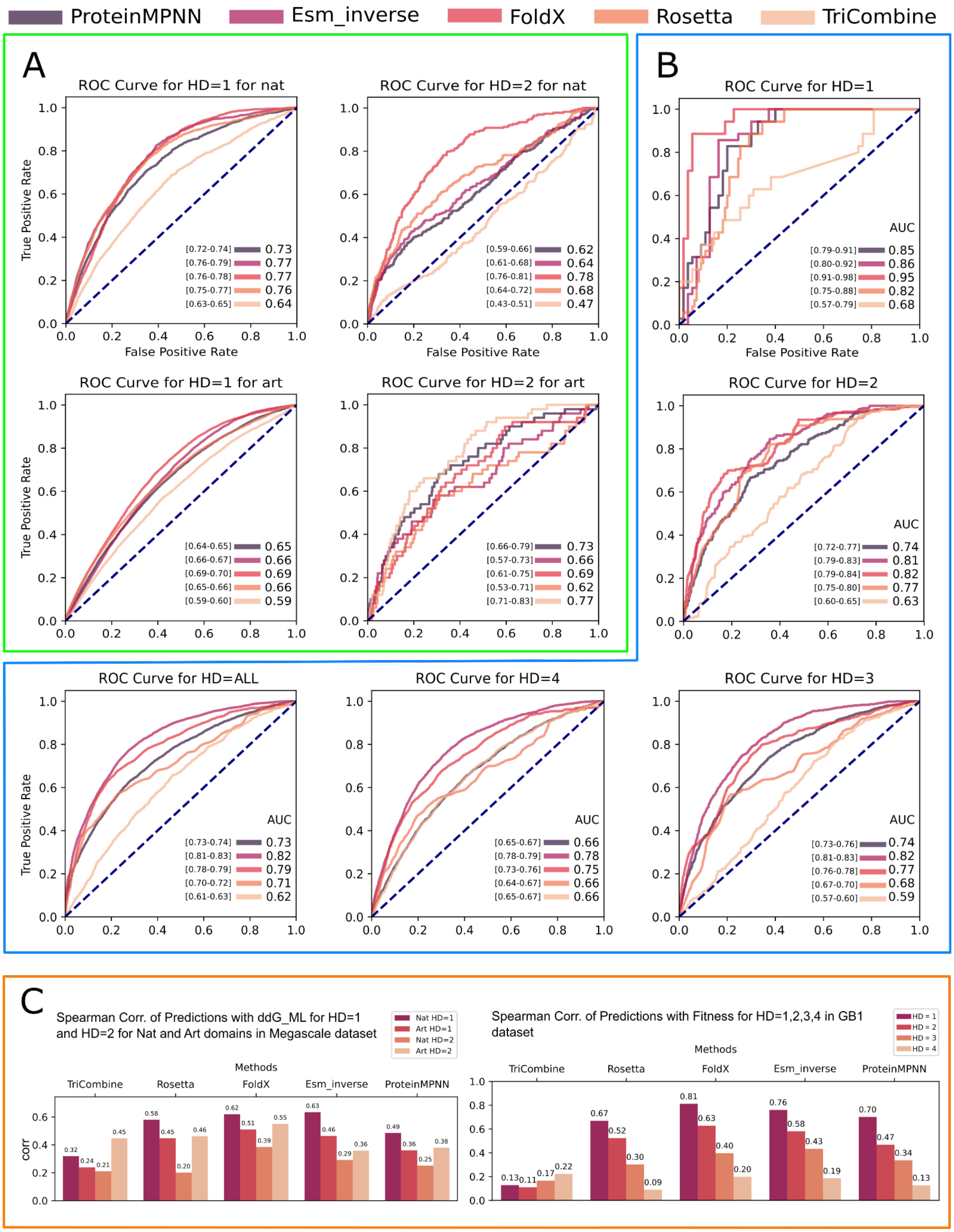


**Supplementary Figure 7: GB1 and mega-scale ROC curves and correlations for predictors performances with respect to experimental stability data**

Supplementary Figure 7A shows ROC curves and AUROC values, including bootstrapped confidence intervals (in square brackets), for FoldX, Rosetta, ProteinMPNN, Esm_inverse, and TriCombine in predicting variants that are more stable than their corresponding wild types in the Megascale dataset. The plots display performance for single (HD=1) and double (HD=2) mutations across both natural and artificial domains enclosed in a green box. Similarly, Figure 7B shows ROC curves and AUROC values for the same tools using the GB1 dataset, enclosed in a blue box. Figure 7C shows Spearman correlations for GB1 and Megascale datasets for the four predictors, in the case of GB1 the four HDs are represented, for Megascale dataset 1 and 2 mutations for natural and artificial domains are showed.

A

hAChE (PDB id: 1hzy)

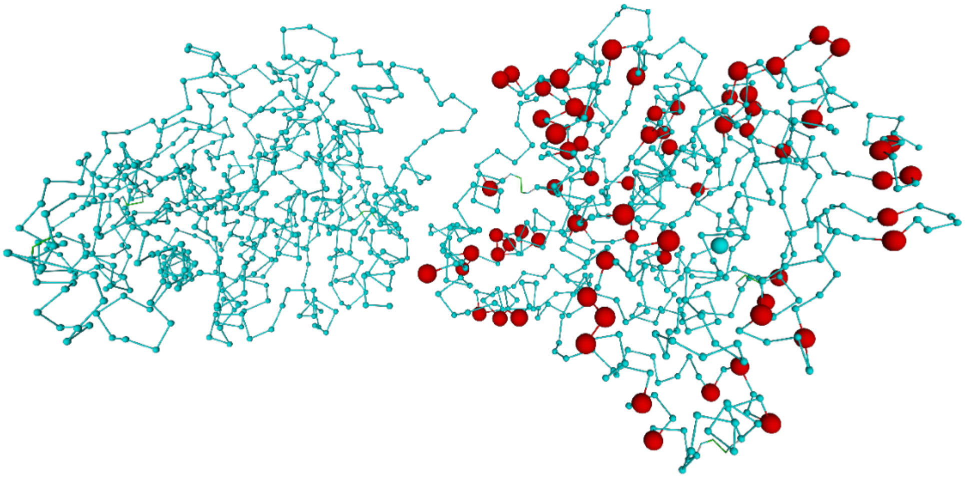

B

TNfn3 (PDB id: 1hzy)

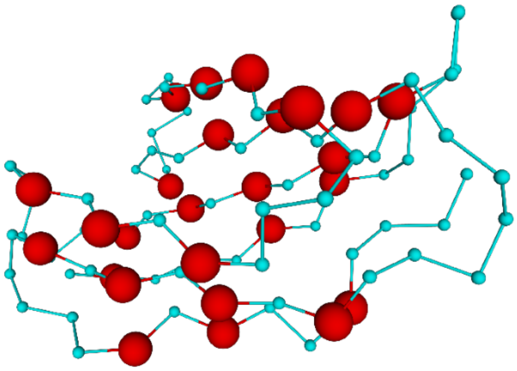

C

PTE (PDB id: 1hzy)

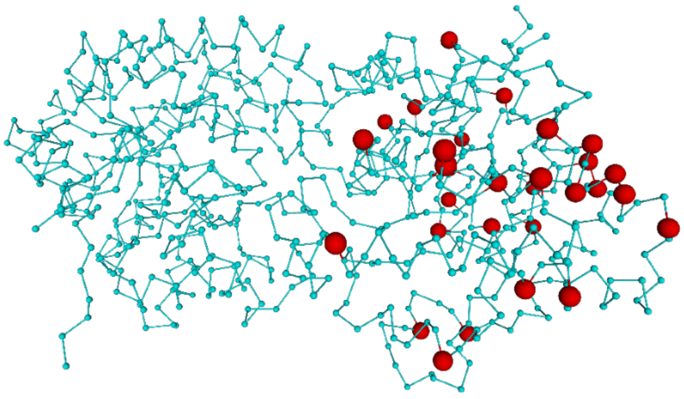

**Supplementary Figure 8: Panoramic of mutated residues for the "bulk" dataset proteins**
Figure shows indiscriminately the alpha carbon atoms (in red spheres) of all mutated residues along all variants for each of the 3 proteins included in this dataset (A: hAChE, B: TNfn3, C: PTE). Proteins are shown as cyan backbones connecting carbon alpha atoms.

### Supplementary tables

**Supplementary Table 1: FoldX sequence details**
The table displays the FoldX output for the command SequenceDetail of all 16 redesigned SH3 cores and the two template structures (1shg, 1e6g). The output has been restricted to the nine designed residues and a column named "mutant diff" shows the total energy calculated by FoldX for the corresponding protein variant. The last four columns show the computed ΔΔGs for the three most important energy components calculated by subtracting to the mutant energy the reference structure ones. mutant diff denotes the FoldX energy difference for the whole protein against its reference template (1e6g for mt2 - mt9 and 1shg for mt11 - mt18); Solv P, the polar solvation; Solv H, the hydrophobic solvation; and VdW clash, van der Waals clashing. Cells are colored from white to red when values are above zero, and red when energies are above FoldX sensitivity (0.8 kcal/mol). Additionally, the energy difference between mutants and their relative starting structures are calculated. The legend under the table explains the FoldX energy terms.

**Supplementary Table 2: Similarity of SH3 variants with the SH3 Pfam family**
The highest number of core amino acid identities shared between a mutant (first column) and any element in the SH3_1 Pfam family (max similarity), along with the count of this value (occurrences), are shown.

**Supplementary Table 3: GDT scores for SH3 mutant structures VS modelling methods for hard (0.5Å) and soft (1Å) thresholds.**
The GDT scores obtained after superposition of the crystallographic structure of the 11 mutants for which a structure was obtained, with respect to each of the modelled structures generated by our comparative analysis. GDTs were measured respectively with thresholds of 0.5Å and 1.0Å.

**Supplementary Table 4: All and C alpha atom RMSD (Å) for all crystallized SH3 variants VS models from various predictors**
The table shows RMSDs for carbon alpha and all atoms of crystallized SH3 variants (1e6g, 1e6h, 1h8k, 1shg, mt12, mt13, mt15, mt16, mt17, mt18, mt8) with respect to the modelled structures of the same mutants, generated with the explored algorithms and combinations (Esm_r_R, Esm_r_F, Esm, FoldX, AlphaFold2_r_R, AlphaFold2_r_F, AlphaFold2, Rosetta, RoseTTAFold_r_R, RoseTTAFold_r_F, RoseTTAFold).

**Supplementary Table 4: Experimental and predicted ΔGs for 36 unique SH3 variants**
The variant stability measurements of 38 SH3, of which 36 are unique mutants and two are experimental stabilities for variants used as design templates (1SHG and 1E6G), as determined

in this work and previously(8). The FoldX predictions and AF2 pLDDT scores for all variants are also shown. A legend explaining FoldX energy terms is available below the table.

**Supplementary Table 5: Experimental and predicted ΔΔGs for 36 unique SH3 variants**

Table shows experimentally determined unfolding stabilities for 36 SH3 variants, the corresponding values produced by the predictors explored in this work and their combinations as ΔΔ-scores, computed by subtracting the predicted stability of the wild type for each of the two batches. The last column labels the SH3 variants in those generated and measured in the current work ("Cianferoni") and those obtained from literature ("Ventura") (8).

**Supplementary Table 6: Mutants crystallization conditions**

**Supplementary Table 7: Rebuilding and refinement data of mutant structures**

**Supplementary Table 8: Experimental and predicted stabilities for hAChE, PTE and TNfn3 variants.**

Table recapitulates the selected variants for the "bulk" dataset. Variants and their stabilities were coarsely inferred from their inactivation (PTE(9)) and melting (hAChE(9), tNfn3(10)) temperatures as reported in the original manuscripts. The Gibbs free energy variation was inferred using the simplified formula:

$$\Delta G(T) = \Delta H_m * (1 - \frac{T}{T_m})$$

Where ΔG(T) is the free energy change at room temperature T, ΔHm is enthalpy change at the melting temperature Tm, T is room temperature, Tm the melting temperature of the protein. The simplified formula lacks the heat capacity change parameter that would allow to compare ΔGs between the three groups of stabilities, which was not available in the original studies. For this reason the stability values and their relationships with the predictions made in our analysis are related to ΔΔG values with respect to the relative wild type variants and are not considered in a correlation analysis putting them together. Variants with melting or inactivation unavailable because unfolded were assigned with a high ΔG value of 6 kcal/mol that made them result as less stable than the wild type in the analysis.


1. J. Mistry, *et al.*, Pfam: The protein families database in 2021. *Nucleic Acids Res.* **49**, D412–D419 (2021).

2. E. Garriga, *et al.*, Multiple Sequence Alignment Computation Using the T-Coffee Regressive Algorithm Implementation. *Methods Mol. Biol.* **2231**, 89–97 (2021).

3. S. R. Eddy, Accelerated Profile HMM Searches. *PLoS Comput. Biol.* **7**, e1002195 (2011).

4. A. R. Viguera, J. C. Martínez, V. V. Filimonov, P. L. Mateo, L. Serrano, Thermodynamic and kinetic analysis of the SH3 domain of spectrin shows a two-state folding transition. *Biochemistry* **33**, 2142–2150 (1994).

5. C. N. Pace, Determination and analysis of urea and guanidine hydrochloride denaturation curves. *Methods Enzymol.* **131**, 266–280 (1986).

6. K. Tsuboyama, *et al.*, Mega-scale experimental analysis of protein folding stability in biology and design. *Nature* **620**, 434–444 (2023).

7. J. P. Mower, PREP-Mt: predictive RNA editor for plant mitochondrial genes. *BMC Bioinformatics* **6**, 96 (2005).

8. S. Ventura, *et al.*, Conformational strain in the hydrophobic core and its implications for protein folding and design. *Nat. Struct. Biol.* **9**, 485–493 (2002).

9. A. Goldenzweig, *et al.*, Automated Structure- and Sequence-Based Design of Proteins for High Bacterial Expression and Stability. *Mol. Cell* **63**, 337–346 (2016).

10. D. N. Kim, T. M. Jacobs, B. Kuhlman, Boosting protein stability with the computational design of β-sheet surfaces. *Protein Sci.* **25**, 702–710 (2016).
